# 5-hydroxymethylcytosine mediated active demethylation is required for neuronal differentiation and function

**DOI:** 10.1101/2021.02.10.430698

**Authors:** Elitsa Stoyanova, Michael Riad, Anjana Rao, Nathaniel Heintz

## Abstract

Although high levels of 5-hydroxymethylcytosine (5hmC) accumulate in neurons, it is not known whether 5hmC can serve as an intermediate in DNA demethylation in postmitotic neurons. We report high resolution mapping of DNA methylation and hydroxymethylation, chromatin accessibility, and histone marks in developing postmitotic Purkinje cells (PCs). Our data reveal new relationships between PC transcriptional and epigenetic programs, and identify a class of genes that lose both 5mC and 5hmC during terminal differentiation. Deletion of the 5hmC writers Tet1, Tet2, and Tet3 from postmitotic PCs prevents loss of 5mC and 5hmC in regulatory domains and gene bodies and hinders transcriptional and epigenetic developmental transitions, resulting in hyper-excitability and increased susceptibility to excitotoxic drugs. Our data demonstrate that Tet-mediated active DNA demethylation occurs in vivo, and that acquisition of the precise molecular and electrophysiological properties of adult PCs requires continued oxidation of 5mC to 5hmC during the final phases of differentiation.

## INTRODUCTION

Development of the mammalian brain requires generation of hundreds of millions of neuronal progenitors that differentiate into distinct cell types with refined functional properties. Although morphological and physiological maturation of most neurons occurs between mid-gestation and a few months or years after birth, the vast majority of CNS neurons must maintain a stable differentiated state for the life of the organism and remain sufficiently plastic to participate in novel behaviors. Studies of signaling molecules and transcriptional programs have identified many mechanisms that orchestrate critical steps in neurogenesis, cell type diversification, neuronal migration, axonal pathfinding and differentiation. Although it has been established that epigenetic regulatory mechanisms are critical for proper development of all cell types, our knowledge of the precise roles of these mechanisms in neuronal differentiation, function and vitality remains rudimentary.

5-hydroxymethylcytosine (5hmC) is produced from 5-methylcytosine (5mC) by the Ten-eleven translocation dioxygenases (Tet1, Tet2, Tet3) (Iyer et al., 2009; Tahiliani et al., 2009). It is present at approximately ten-fold higher levels in neurons than peripheral cell types (Globisch et al., 2010; Kriaucionis and Heintz, 2009), and its distribution across the genome of adult neurons is cell specific and correlated with active gene expression (Mellén et al., 2017, 2012; Szulwach et al., 2011). Stable accumulation of 5hmC in neurons contributes to the function of the essential neuronal protein MeCP2 (Chen et al., 2015, p. 2; Mellén et al., 2012, p. 2). Thus, the genomic occupancy of MeCP2 reflects its differential affinities for methylated and hydroxymethylated CG and CH dinucleotides (Ayata, 2013; Gabel et al., 2015; Mellén et al., 2017), and transcriptional alterations in MeCP2 knockout mice are correlated with the levels of 5mC and 5hmC in expressed genes (Mellen et al, 2017). Imaging of the binding and diffusion of single MeCP2 molecules in living neurons lacking Dnmt3a or Tet 1,2, and 3 demonstrate that its binding is exquisitely sensitive to the levels of 5mC and 5hmC (Piccolo et al., 2019). Given these data and the sensitivity of neuronal function to MeCP2 gene dosage (Chahrour et al., 2008; Nan and Bird, 2001), relief of the repressive functions of MeCP2 through Tet-mediated conversion of high affinity 5mCG binding sites to low affinity 5hmCG sites provides an important mechanism for modulation of chromatin structure and transcription.

In dividing cells, 5hmC serves as an intermediate in DNA demethylation because maintenance DNA methyltransferases do not recognize hemi-hydroxymethylated cytosines in order to reestablish methylation (Wu and Zhang, 2017). Consequently, 5hmC is lost passively and replaced by C as a result of replicative dilution. Loss of function studies in mouse embryonic stem cells (ESCs) (Dawlaty et al., 2013) and lymphocyte lineages (Lio and Rao, 2019) have demonstrated that the role of Tet mediated replicative demethylation is to provide full accessibility to regulatory regions necessary for expression of genes required for differentiation. For example, at the activation-induced deaminase (AID) locus in B cells, Tet activity is required for demethylation and activation of enhancer regions that modulate AID expression to enable class switch recombination (Lio et al., 2019). Although Tet mediated replication dependent passive demethylation is thought to be common in many dividing cell types, including progenitor cells in the developing forebrain (Rudenko et al., 2013), hippocampus (Guo et al., 2011, p. 1) and cerebellum (Zhu et al., 2016), chromatin remodeling by this pathway cannot contribute to programs responsible for the dramatic increases in size, morphological complexity and connectivity that occur during the post-mitotic differentiation of most projection neurons.

A second pathway for DNA demethylation that does not depend on DNA replication, often referred to as active demethylation, is based on the finding that 5-hydroxymethylcytosine can be further oxidized by Tet proteins to produce 5-formylcytosine (5fC) and 5-carboxylcytosine (5caC). Removal of 5fC and 5caC by thymine DNA glycosylase (TDG) dependent base excision repair (BER) provides another mechanism for 5hmC mediated DNA demethylation (He et al., 2011). Clear evidence that this pathway can operate in cultured cells has been presented, and detailed biochemical studies have delineated the mechanisms operating in TDG-BER DNA demethylation (Weber et al., 2016). Although active DNA demethylation is ideally suited for remodeling DNA methylation in post-mitotic neurons, in vivo evidence demonstrating a role for this third function of 5hmC in neuronal development or function is lacking.

Here, we report single nucleotide resolution studies of DNA methylation and hydroxymethylation, transcription, chromatin accessibility, and H3K4me3 and H3K27me3 histone marks in post-mitotic, differentiating Purkinje cell. As Purkinje cells transition from relatively small, multipolar immature cells to fully elaborated large neurons with complex dendritic arbors and hundreds of thousands of synapses, DNA methylation and hydroxymethylation are reconfigured as epigenetic and transcriptional programs progress. Our data confirm previous studies of the relationships between transcription, DNA methylation and chromatin organization (Lister et al., 2013; Mellén et al., 2017; Tsagaratou et al., 2017, 2014), and they reveal several novel modes of transcriptional activation and repression. Notably, we identify a class developmentally induced Purkinje cell specific genes that are highly expressed and demethylated in the final stages of Purkinje cell differentiation. Studies of a newly generated Tet1, Tet2, Tet3 Purkinje cell specific triple knockout (Pcp2CreTetTKO) mouse lines demonstrate that transcriptional and epigenetic maturation in Purkinje cells, including active demethylation of these late expressed genes, requires continued oxidation of 5mC to 5hmC. Furthermore, electrophysiological recordings from Pcp2CreTetTKO Purkinje cells shows that loss of Tet activity in the second postnatal week results in hyperactivity. Finally, Pcp2CreTetTKO mice display enhanced sensitivity to the excitotoxic drug harmaline. These data demonstrate definitively that active demethylation occurs in postmitotic neurons, and that 5hmC plays an essential role in late stages of neuronal differentiation.

## RESULTS

In mice, Purkinje cell progenitors complete their final cell cycle between e10.5 and e13.5 within the ventricular zone of the developing cerebellar anlage (Leto et al., 2016). As they exit cell cvcle and commit to a Purkinje cell fate, they express the transcription factors Lhx1/Lhx5 (Chizhikov, 2006; Morales and Hatten, 2006). To characterize 5hmC accumulation, transcription and chromatin organization during their postmitotic development, we chose to analyze PCs at the start of their accelerated differentiation (P0), during their rapid phase of morphological development (P7) and as fully mature, differentiated neurons (adult) (Figure 1A-B). We employed fluorescence activated nuclear sorting (FANS) to purify Itpr1 positive PC nuclei in two biological replicates (Figure 1C-D, Figure S1A) (Xu et al., 2018). Since PCs are extremely rare and difficult to isolate, we optimized low input protocols for genomic profiling. We used ~20,000 nuclei for transcriptional profiling, ~200,000 nuclei for bisulfite sequencing (BS-seq) and oxidative bisulfite sequencing (oxBSseq), 25,000 nuclei for the assay for transposase-accessible chromatin sequencing (ATACSeq), and 25,000 nuclei for H3K27me3 and H3K4me3 chromatin immunoprecipitation sequencing (ChIPSeq) (Figure 1E-F). Analysis of biological replicates for each of the assays revealed that the sorted nuclei expressed markers of Purkinje neurons, and did not express genes marking the two most abundant cerebellar cell types, granule cells or glia (Figure S1B). The Pearson correlation coefficients between each replicate at all timepoints and techniques (RNAseq, OxBSSeq, ATACSeq and ChIPSeq) were high (over 0.9), showing strongly reproducible results (Figure S1C-F). Bisulfite conversion and oxidation rates for the OxBSSeq datasets were within the expected range (Figure S1G).

**Fig. 1.**
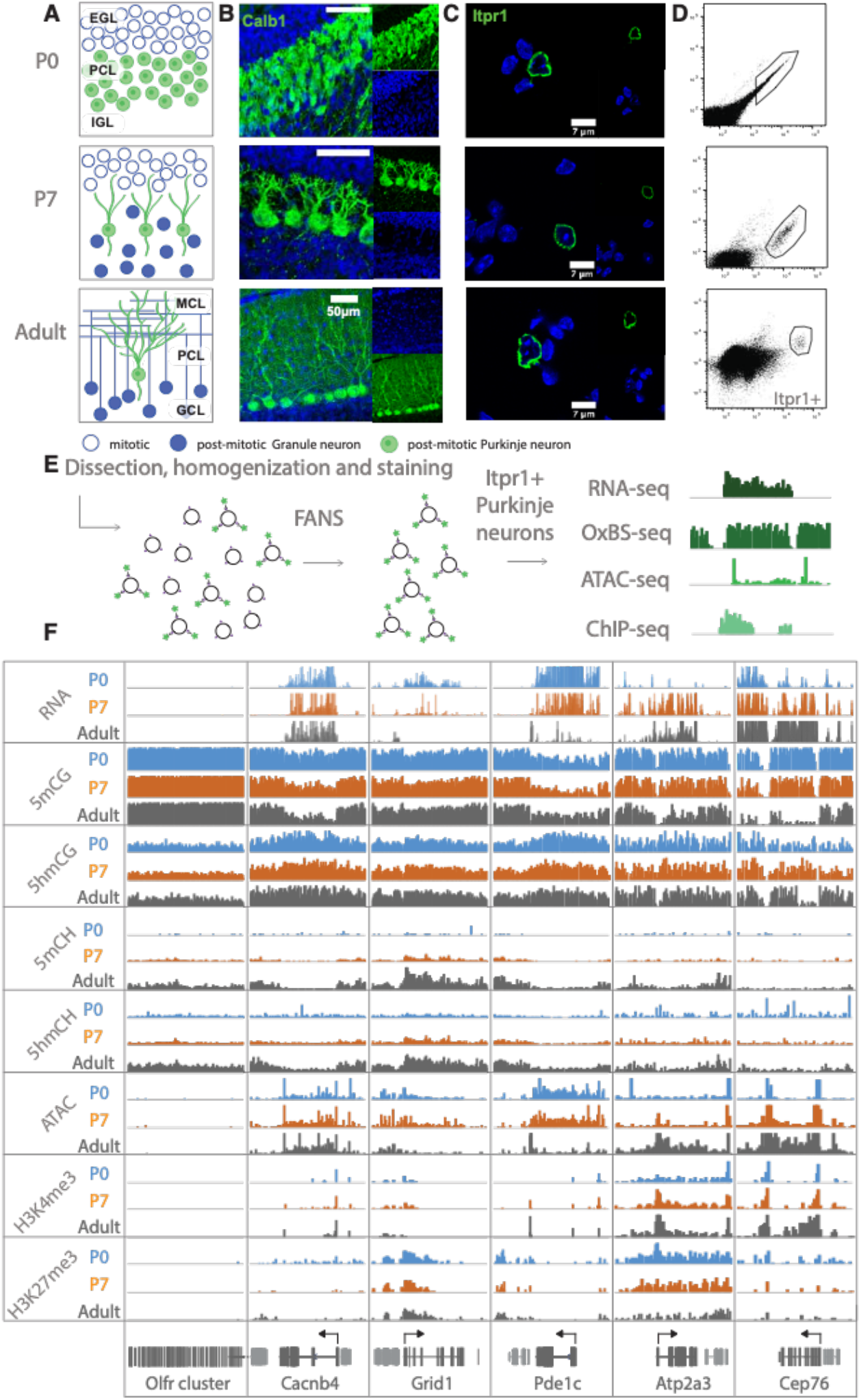
Chromatin landscape in differentiating Purkinje neurons. **A**. Schematic of PC differentiation and growth at P0, P7 and adult (approximately 8 weeks old) timepoints. EGL-external granule layer, PCL-Purkinje cell layer, IGL-internal granule layer, MCL-molecular cell layer, GCL-granule cell layer. **B**. Immunofluorescence staining with Calb1 (green) of PCs in murine cerebella at P0, P7 and adult timepoints. **C**. Example of Purkinje cell nuclei stained with Itpr1 (green) post dissociation and pre-sorting, counterstained with DAPI, a heterochromatin marker. **D**. Representative plots of fluorescence activated nuclear sorting of Purkinje cells at P0, P7 and adult timepoints with Itpr1. **E**. Workflow schematic of nuclei isolation, antibody staining with anti-Itpr1, fluorescence activated sorting, and downstream sequencing applications. **D**. Integrated genome viewer (IGV) representation of example regions of differentially regulated genes (Olfr cluster – always silent, Cacnb4 – always expressed, Grid1 and Pde1c – developmentally downregulated, Atp2a3 and Cep76 – developmentally upregulated). Top tracks show RNA expression in RPKM (reads per kilobase per million mapped reads), mCG tracks show methylation level in CG context from 0 to 0.8, hmCG tracks show hydroxymethylation level in CG context from 0 to 0.45, mCH and hmCH show methylation and hydroxymethylation level in CH context (H = A, C or T) from 0 to 0.04. ATAC tracks show ATAC-seq read density in RPKM from 0 to 10. H3K4me3 tracks show input normalized enrichment in RPKM from 0 to 5. H3K27me3 tracks show input normalized enrichment in RPKM from 0 to 3.

It is important to note that we chose to focus on OxBSSeq (Booth et al., 2013) as the most definitive technology for analysis of DNA methylation because, in contrast to BS-Seq alone, it allows us to assess the contributions of both 5-methylcytosine and 5-hydroxymethylcytosine to neuronal development. Given the distinct functions of these two modifications, analysis of their separate contributions to epigenetic regulation of the genome is more informative than BSSeq alone. For example, in many genes, 5mCG is depleted and 5hmCG accumulates in the gene body as its expression increases between P0 and adult. This is clearly evident for the gene encoding the cyclic GMP-dependent protein kinase (Prkg1), whose expression in PCs has been shown to be essential for long term depression (Feil et al., 2003). In this case, the OxBSSeq data reveal a transition from 5mCG to 5hmCG that cannot be detected in the BS-Seq data as the gene in activated between p0 and adult (Figure S1H). These data illustrate that a precise evaluation of DNA methylation status in relation to other programs unfolding during development for this gene and many others is advanced significantly by inclusion of OxBSSeq data.

### Transcriptional programs altered during Purkinje cell differentiation

Purkinje cell progenitors complete their final divisions in the cerebellar primordium between e11-e13 (Butts et al., 2014) and remain as a multilayered, simple migrating cell population until birth (P0). In the first postnatal week, they organize into a monolayer as the cerebellum enlarges and begin the transition from a multipolar primitive neuronal morphology to one of the largest neurons in the brain with a characteristic, highly elaborate planar dendritic arbor. As they begin the second postnatal week (P7), Purkinje cells undergo an important developmental transition that includes refinement of their climbing fiber input, formation of many thousands of parallel fiber synapses, and myelination of their axons (Leto et al., 2016). The tremendous increase in synaptogenesis and connectivity continues until PCs attain their mature morphology and functions at 3 to 4 weeks of age. As these programs unfold, there is a global decrease of 5mCG and global increase of 5hmCG (Figure S2A).

To identify genes whose transcription is stable and those that are dynamically regulated during PC differentiation, we conducted differential expression analysis between P0 and adult Purkinje neurons, filtering for significance of p<0.01 and log2 fold change of > 2 in either direction (Figure 2A-B,). Using these criteria, there were 922 developmentally repressed genes (down regulated in adult), enriched in categories involved in cell signaling and axon guidance pathways (Grid1, Pde1c Figure 1F). We identified 432 genes with increased expression as PCs differentiate, encoding proteins known to be important for the mature functions of PCs including calcium ion buffering and transport, regulation of the inositol triphosphate signaling pathway and RNA splicing. As expected, similar analysis comparing P0 and P7 or P7 and adult data revealed genes overlapping with those identified in the overall P0 and adult comparative data (Figure S2B-D), although additional categories of RNA metabolism, synaptic signaling and ion transport are enriched in the P7 to adult analysis (Figure S2C). As anticipated, the P7 profiles we have included in our analysis have been essential in assessing both the developmental course of epigenetic events studied here and the consequences of loss of 5hmC in PCs (see below).

**Fig. 2.**
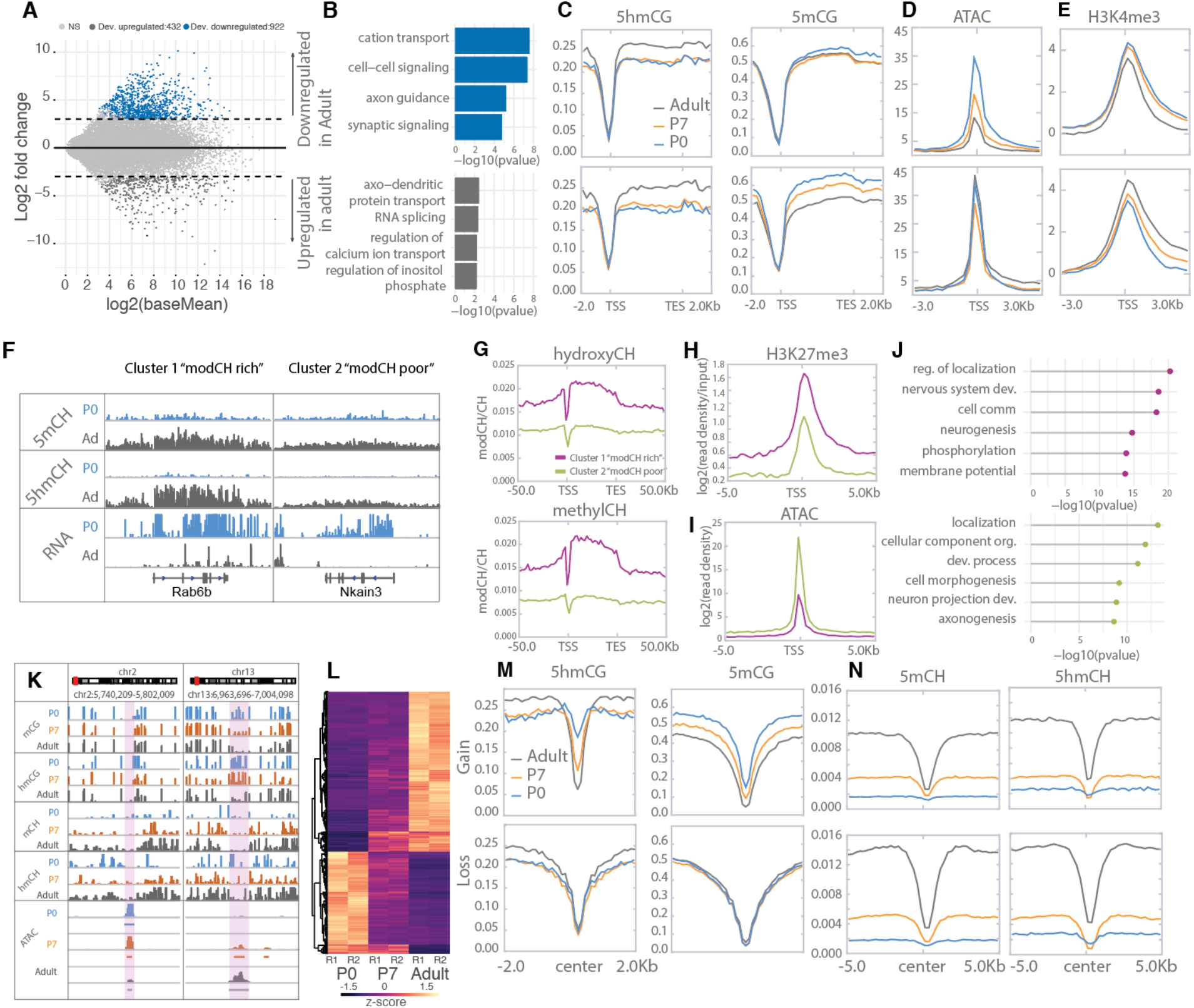
Cytosine modification dynamics in in relation to differential expression and accessibility. **A**. MA plot representing statistically significant (p < 0.01, log2(fold change) > 2) differential gene expression between P0 (blue) and adult (dark gray). Light grey dots represent genes that are not statistically significant. **B**. Gene ontology analysis of the developmentally up- and down-regulated genes. **C**-**E**. Metagene plots representing the mean value of 5hmCG/CG, 5mCG/CG (**C**), ATACSeq read density (**D**) and H3K4me3 accumulation (**E**) over the gene bodies and promoters of developmentally up- and down-regulated genes over the three timepoints (P0 – blue, P7 – orange and adult – dark grey). **F**. Genome browser representation of the two classes of developmentally down-regulated genes with differential accumulation of CpH modifications. **G-I**. Metagene plots representing the mean value of 5hmCH/CH, 5mCH/CH (**G**), ATACSeq read density (**H**) and H3K27me3 accumulation (**I**) over the gene bodies and promoters of developmentally down-regulated genes divided into two clusters by k-means clustering analysis (green for “modCH poor”, purple for “modCH rich”) **J**. Gene ontology analysis of the two clusters. **K**. Genome browser representation of two regions with differential accessibility. **L**. Heatmap representing the differentially accessible regions between P0 and adult PCs (p < 0.01, log2(fold change) > 4). **M**-**N**. Metagene plots representing the mean values of 5hmCG/CG and 5mCG/CG (**M**) and 5hmCH/CH and 5mCH/CH (**N**) over the centers and flanking regions of peaks that gained or lost accessibility relative to adult PCs.

### Constitutively expressed genes are epigenetically stable

Despite the many genes that are dynamically regulated during these developmental transitions, the majority of genes in PCs are either silent or constitutively expressed. Approximately half of the genes in the mouse genome (e.g genes encoding olfactory receptors (Figure 1F), immunoglobulins, hemoglobin subunits, etc) are never expressed in PCs and remain heavily methylated and inaccessible. As previously reported for other neurons (Mo et al., 2015), there is a second class of silent genes in PCs that are enriched in transcription factors expressed in very early embryos (Hox clusters, other homeobox genes, etc). They are completely demethylated and enriched for the H3K27me3 histone mark, indicating that they are repressed by the Polycomb Repressive Complex (Li et al., 2018).

Actively transcribed genes whose levels vary little during PC differentiation, for example the calcium channel auxiliary subunit Cacnb4 (Figure 1F), are also epigenetically stable. This is a large (~3000) and diverse class of genes that carry previously characterized activating epigenetic marks: low levels of 5mCG, 5mCH, and 5hmCH; elevated levels of 5hmCG; they are ATAC accessible; and their promoters carry the activating histone mark H3K4me3. As expected, the levels of these marks vary widely between genes and generally reflect the level of expression. As PCs mature the conversion of 5mC to 5hmC continues to increase within the gene bodies of the most active of these genes without a strong impact on expression.

### Epigenetic signatures reveal two classes of developmentally repressed genes in PCs

To identify mechanisms associated with developmental repression, we analyzed features previously associated negatively with transcription in the 922 genes whose expression decreases during PC differentiation. A consistent finding is that in most genes decreased expression is associated with a loss of promoter accessibility as assayed by ATACseq (Figure 2D, Figure S2D). These changes are not correlated with changes in DNA methylation or H3K27me3 occupancy (Figure 2C,D). Furthermore, 5hmCG accumulated over the genes bodies early in development is not removed (Figure 2C), suggesting that the presence 5hmCG is not sufficient to maintain transcription. Interestingly, analysis of the repressive marks 5mCH and 5hmCH reveals at least two distinctly recognizable epigenetic patterns. Repressed genes that accumulate 5mCH and 5hmCH over their gene bodies (modCHrich) are associated with lower promoter accessibility and enhanced accumulation of H3K27me3 relative to those repressed genes with low levels of CH modification (modCHpoor) (Fig. 2F-J, FigureS2F-G.). Although the data for modCHrich genes is consistent with transcriptional inhibition through both polycomb repressive complexes and MeCP2, our data do not identify the mechanisms responsible for repression of the modCHpoor genes.

### Loss of DNA methylation in a subset of developmentally activated PC expressed genes

We identified 432 genes that increase in expression as PCs differentiate. In general, these genes encode proteins known to be important for the mature functions of PCs, including calcium ion buffering and transport, regulation of the inositol triphosphate signaling pathway and RNA splicing. The chromatin landscape of genes whose expression increases during Purkinje cell differentiation is similar to those that are constitutively transcribed (e.g. Atp2a3, Cep76 Figure 1G). Their promoters become highly accessible, they accumulate high levels of H3K4me3, their gene bodies progressively lose 5mCG and gain 5hmCG, and they do not accumulate 5mCH or 5hmCH. Although there are many precedents for acquisition of these activating features as genes become expressed, our data reveal an additional, surprising feature of gene activation that has not been documented in postmitotic cells. Thus, in a small subset of highly expressed genes (e.g. Cep76, Itpr1, Mtss1) we noticed a profound loss of both 5hmCG and 5mCG during the terminal, postmitotic stage of PC differentiation (Fig. 1D, 2). To investigate the apparent demethylation over this class of genes, we computationally identified DNA methylation valleys (DMVs) as previously described (Jeong et al., 2014; Mo et al., 2015; Xie et al., 2013). In brief, DMVs were identified by filtering undermethylated regions (UMRs) for length (>5kb) and merging any regions within 1kb (Burger et al. 2013) (Figure S3A-B, Table S1). We then divided the DMVs into accessible or inaccessible based on their level of ATAC-Seq read enrichment (Figure S3C). When exploring the length of the DMVs, we noticed multiple regions longer than 15kb, which was very interesting as the average length of active DMVs is about the size of a promoter (~5kb) (Figure S3D) (Jeong et al. 2014). Furthermore, we show that DMVs are correlated with the presence of extremely broad ATAC and H3K4me3 peaks (some up to 100kb) (FigureS3E). Broad DMVs (>15kb) were present over both active and inactive genes, including the previously defined class of early developmental transcription factors that are inactive, hypo-methylated, inaccessible, and have elevated H3K27me3 levels (Figure 3A, Foxd1). The evolution of DMVs in differentiating PCs is complex, and it reflects the transcriptional status of individual genes. Itpr1 and Mtss1 transcription steadily increases from P7 to adult, and it is accompanied by decreases in the level of 5mGC levels over their gene bodies. Both genes have high levels of 5hmCG at P0 which steadily decrease during differentiation. Analysis of the BS-Seq data (5hmCG and 5mCG combined) shows multiple hypo-DMR (differentially methylated regions) present at the adult timepoint compared to P0 (Figure 3A, hypoDMR track). As methylation and hydroxymethylation are lost and DMVs broaden (FigureS3 I-K), chromatin accessibility and H3K4me3 accumulation increase correspondingly (Figure 3B-E).

**Fig. 3.**
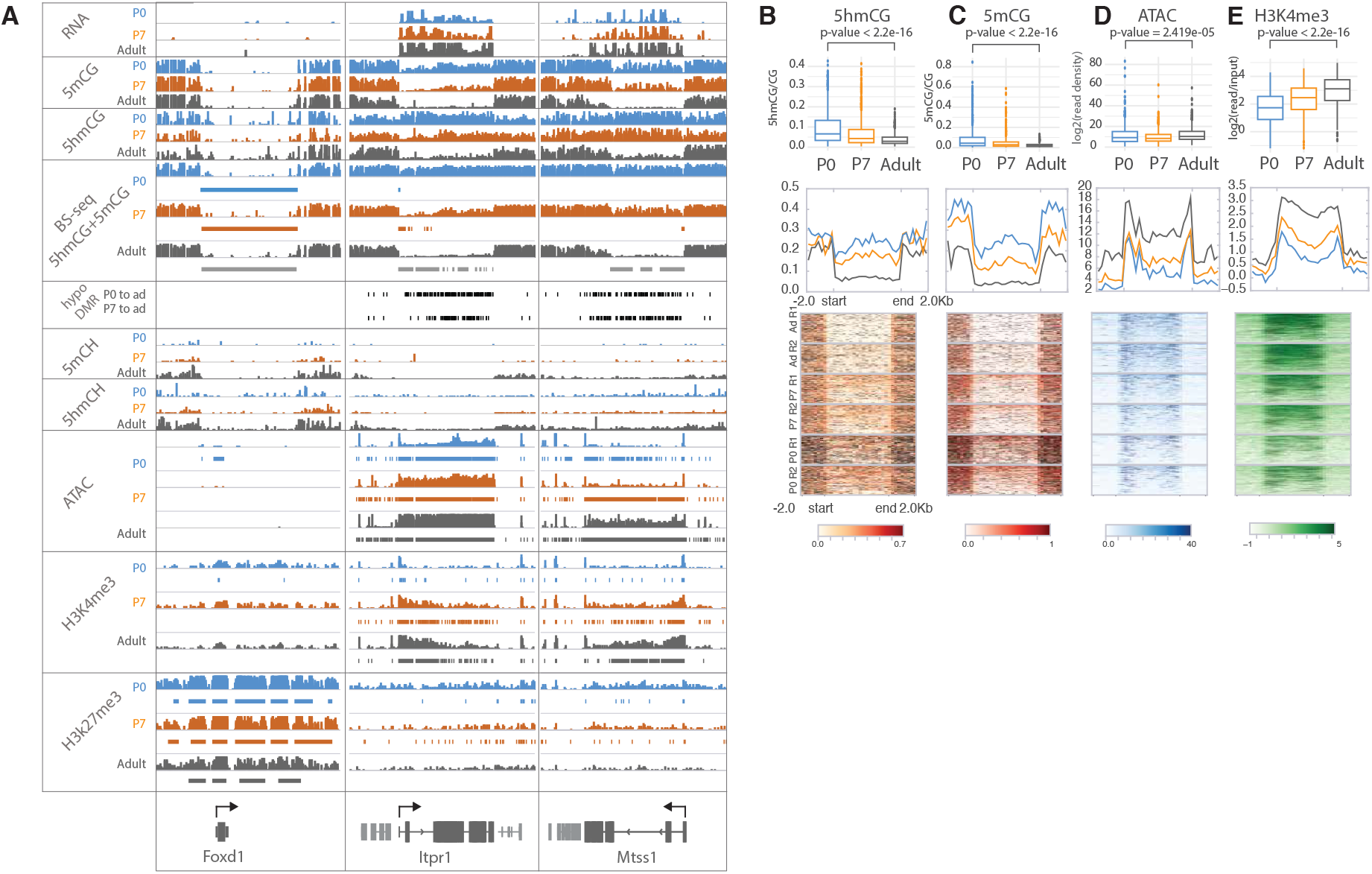
Loss of 5mCG and 5hmCG in a novel class of epigenetically regulated highly expressed Purkinje specific genes. **A**. IGV representation of example regions of inactive DMV (Foxd1) and active PC specific DMVs (Itpr1, Mtss1). BSSeq tracks shows combined levels of mCG and hmCG ranging from 0 to 1. Bars under BSSeq tracks are computationally identified DMVs, black bars denote hypo-DMRs in adult, compared to P0 or P7. Bars under ATAC-Seq, H3K4me3 and H3K27me3 tracks denote broad peaks of signal enrichment. **B**-**E**. 5hmCG/CG (**B**), 5mCG/CG (**C**), ATACSeq log2(read density) RPKM enrichment (**D**) and H3K4me3 log2(input normalized) RPKM enrichment (**E**) quantification over DMVs at each timepoint. Boxplots show mean value per DMVs, the test for significance is Wilcoxon. Metagene plots show mean value ± 2kb around the DMV region, regardless of gene directionality. Heatmaps show data summarized in the metagene plots.

Interestingly, many of the genes associated with active broad DMVs are PC specific and very highly expressed (Figure S3F). Gene ontology analysis indicates that they are involved in the inositol triphosphate/calcium signaling pathways, and a subset is associated with ataxia and autism (Figure S3H). Neither the accessible nor inaccessible DMVs accumulated modified cytosines in CpH context (Figure S3G). These data provide strong evidence that DNA demethylation can occur in postmitotic neurons, and that it is enhanced in a specific class of genes that are extremely highly expressed and functionally important. Although mass spectroscopic analysis of genomic DNA from differentiating PCs (Figure S3L) failed to detect 5-formylcytosine or 5-carboxycytosine, their involvement as transient intermediates in the loss of 5mC and 5hmC cannot be ruled out because the small fraction of the genome covered by this gene class may preclude their detection (Guo et al., 2011; He et al., 2011; Ito et al., 2011).

Previous studies of enhancers and other small regulatory sites have established that their activation is accompanied by increased ATAC accessibility and loss of DNA methylation (Lio et al., 2016). To determine whether loss of DNA methylation can occur also in putative enhancers as well as large DMRs (as documented above), we used an established computational method (Burger et al., 2013) to identify small regions with statistically significant ATACSeq signal enrichment (Figure 2 K-N). We then employed differential accessibility analysis to divide those regions into two groups based the magnitude (log2 fold change > 4) and significance (p<0.01) of their changes during Purkinje cell differentiation. Those that became more accessible between P0 and adult (log2 fold change > 4) experienced a “gain” and those whose accessibility diminished during this time are characterized by a “loss”. As anticipated, those ATAC peaks that gain accessibility lose both 5hmC and 5mC as cells progress from P0 to adult (Figure 2N, Gain section), whereas no significant changes in methylation at ATAC peak centers occur in sites that lose accessibility (Figure 2N, Loss section). This is consistent with our observation that DNA demethylation occurs in a subset of gene bodies during PC differentiation. Although these ATAC peaks have not been identified as active enhancers, it is noteworthy that the transcription factor motifs found in these different classes of ATAC sites are distinct (Figure S2H), and that our data are consistent with prior studies indicating that enhancer activation is accompanied by enhanced ATAC accessibility and DNA demethylation. In this case, however, DNA demethylation does not require cell division.

### Purkinje cell specific Tet1, Tet2, Tet3 triple knockout mouse lines

Despite convincing evidence that Tet-mediated active DNA demethylation can occur in mouse embryonic stem cells through the TDG-BER pathway (He et al., 2011; Weber et al., 2016), evidence that this can occur in vivo has been difficult to obtain because this requires loss or complete inhibition of all three Tet oxidases in a single cell type after cells have exited the cell cycle permanently. To provide this evidence, we generated Purkinje cell specific Tet1/Tet2/Tet3 triple knockout (Pcp2CreTetTKO) mouse lines (Figure 4A). We chose to drive Cre recombinase expression using an engineered Pcp2 BAC employed previously to generate accurate Cre driver lines (Gong et al., 2007) because the onset of expression of the PCP2 gene and the corresponding BAC vector occur approximately one week after birth as the cerebellum enters its terminal phase of development and maturation. The Pcp2Cre BAC was introduced by pronuclear injection into ova from females carrying floxed alleles of all three Tet genes (Figure S4A) yielding several founder lines. PCR analysis of the floxed regions of each Tet gene in purified PC genomic DNA, and RNA-seq analysis confirmed the deletion of exons from all three TET proteins (Figure S4B-C). The two lines chosen for analysis displayed no gross motor phenotype, survival was normal, and recombination activity was specific to Purkinje cells (Figure 4B-D). Cre recombinase activity, and thus recombination in the Tet genes, began at approximately P7 more than two weeks after PCs have completed their last division (Figure S5A-D). The deletion of the Tet proteins did not affect the expression of Itpr1 significantly and we confirmed that it could still be used for purification of Purkinje cells (Figure S4D-E). Evaluation of the quality control metrics of the datasets showed strong correlation coefficients, and enrichment for Purkinje markers (Figure S4F-G). These lines, therefore, allow assessment of the consequences of loss of Tet activity during the rapid phase of cell growth in postmitotic, differentiating Purkinje neurons.

**Fig. 4.**
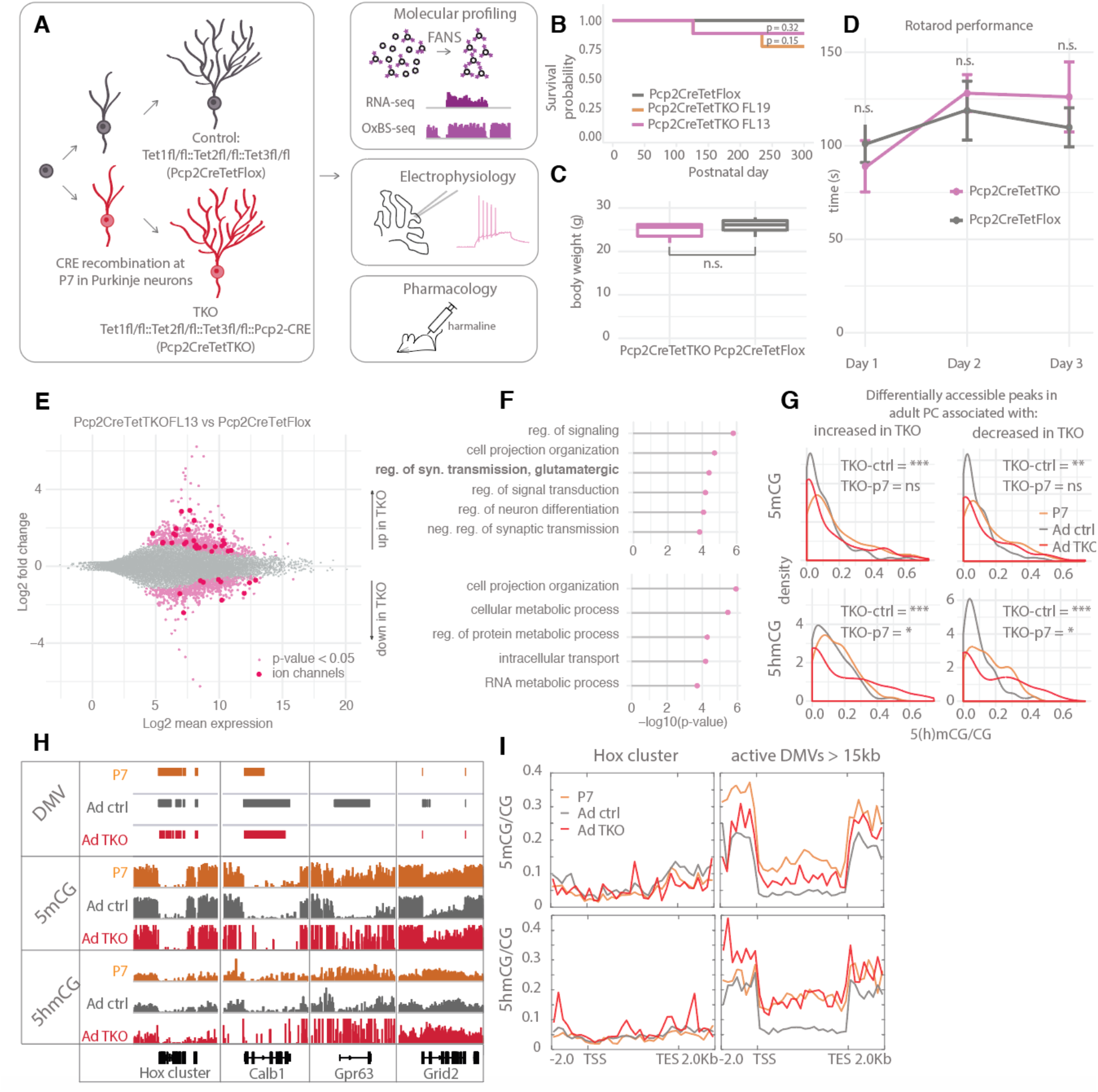
Loss of Tet activity leads to impaired gene expression regulation in PCs. **A**. Schematic of experimental design to probe the triple Tet1,2,3 knock-out effect in adult PCs. **B**. Kaplan-Meier curve representing the survival fraction of Pcp2CreTetFlox (n=8) and Pcp2CreTetTKO (Founder13 n=8, Founder19 n=8) mice since date of birth. **C**. Body weight of Pcp2CreTetFlox and Pcp2CreTetTKO at 8 weeks. **D**. Rotarod evaluation of motor skills in Pcp2CreTetFlox and Pcp2CreTetTKO at 8 weeks. **E**. MA plot representing differential gene expression analysis between Pcp2CreTetFlox and Pcp2CreTetTKO. Pink dots represent genes with p-value < 0.05, red dots - ion channels. **F**. Gene ontology analysis of statistically significant differentially expressed genes. Top panel shows categories of genes with increased expression in Pcp2CreTetTKO, bottom panel shows genes with decreased expression in Pcp2CreTetTKO. **G**. Density plots showing differential levels of 5hmCG/CG and 5mCG/CG over differentially accessible peaks only present in adult PCs. **H**. IGV representation of example DMVs affected by Pcp2CreTetTKO. Hox cluster and Calb1 show minor changes in length as they are established before the P7 onset of Cre. Gpr63 and Grid2 show significant reduction in length as they are established after P7. Solid bars represent DMVs identified at each condition. **I**. Quantification of 5hmCG/CG and 5mCG/CG over the two classes of DMVs – Hox cluster and large active DMVs.

### DNA demethylation in postmitotic Purkinje cells requires 5hmC

Loss of function studies of Tet1, Tet2, Tet3 and combinations thereof in ESCs and lymphocyte lineages have established that 5hmC plays a direct role in transcriptional regulation as a consequence of Tet mediated replication dependent loss of 5mC in enhancers, promoters and gene bodies. To determine whether disruption of ongoing 5hmC accumulation in postmitotic Purkinje cells results in altered transcriptional regulation despite the inability to utilize passive DNA demethylation as a regulatory mechanism, we first compared the transcriptome of adult Pcp2CreTetTKO mice with floxed Cre negative littermates (Pcp2CreTetFlox). We identified genes with both increased and decreased levels of expression, including a large number of ion channels and receptors involved in synaptic transmission (Figure 4E-F). To assess whether this reflects a direct role of TET proteins in transcriptional regulation as a consequence of loss of 5mC in enhancers, promoters and gene bodies, we first identified candidate regulatory domains by comparative analysis of P0 and adult ATACSeq data to detect changes in chromatin accessibility that occur during PC differentiation. We then focused on those regions associated with genes that are transcriptionally impacted in the Pcp2CreTetTKO. Analysis of ATAC accessible regions that are present only in adult PCs revealed that they have significantly more 5mCG and 5hmCG in P0 and P7 PCs than in the adult, and that loss of DNA methylation in these sites does not occur in the Pcp2CreTetTKO (Figure 4G). The modification status of those regions is similar between P7 and Pcp2CreTetTKO, compared to the Pcp2CreTetFlox adult. This is also reflected in the expression levels of the Pcp2CreTetTKO downregulated genes, which are similar to P7 (Figure S5E) Cytosine methylation status over the promoters for these genes is not altered (Figure S5F). Furthermore, genes that acquire broad DMVs late in development over the entire gene (Gpr63) or partially (Grid2), are no longer demethylated in the Pcp2CreTetTKO, whereas DMVs established early in development in either transcriptionally silent (Hox cluster, e.g. Hoxa10) or active (Calb1) genes are not affected (Figure 4H-I, Figure 5SG-H). These data demonstrate that ongoing oxidation of 5mCG to 5hmCG by Tet oxidases in PCs is required for execution of molecular and epigenetic programs associated differentiation, and that DNA demethylation of regulatory domains and gene bodies occurs in PCs the absence of cell division.

**Fig. 5.**
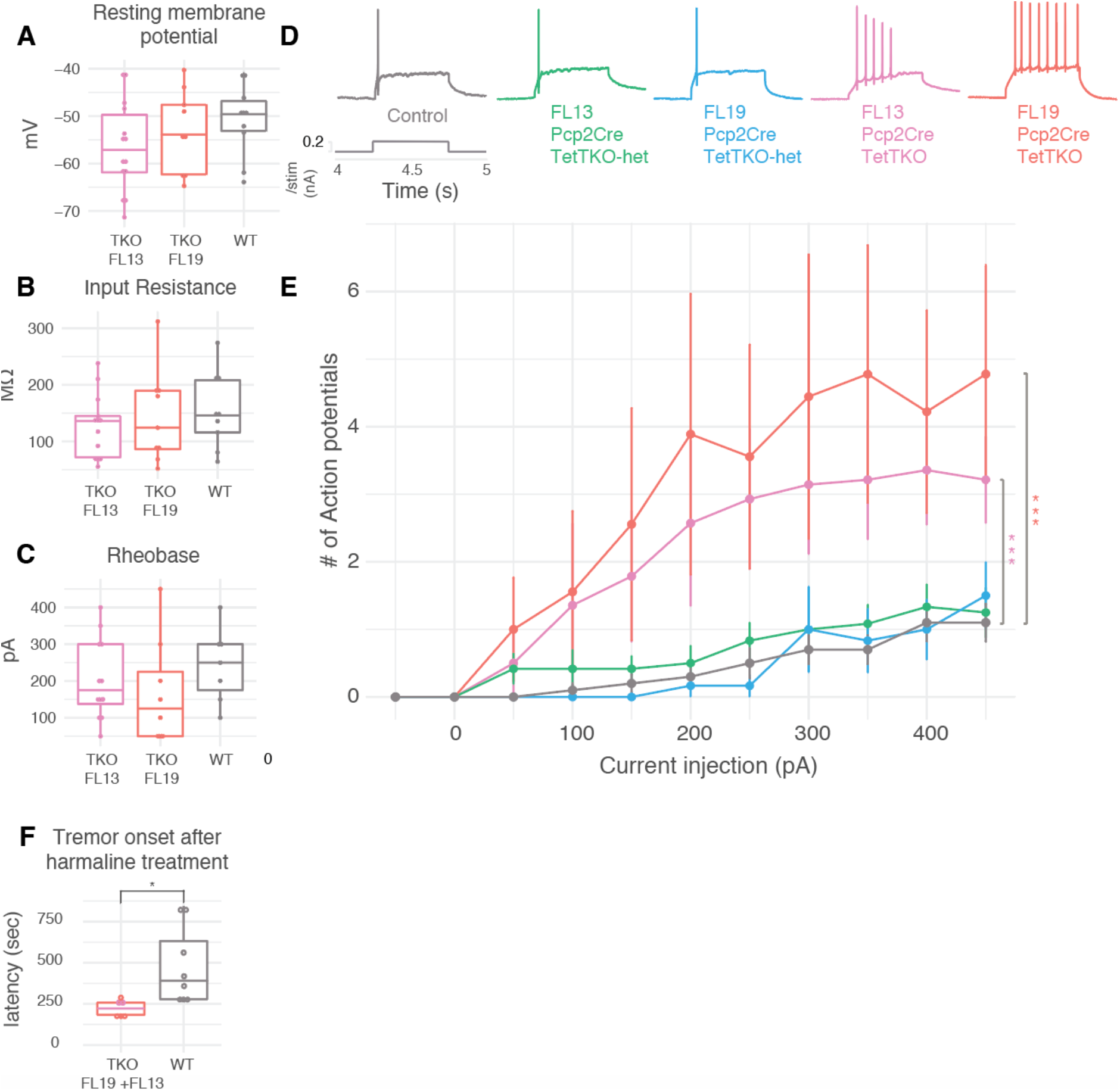
Conditional Tet 1,2,3 deficiency causes hyper-excitability and increased susceptibility to excitotoxic drugs. **A**. Resting membrane potential, recorded after breaking the membrane with no current injected. **B**. Input resistance at −50pA current injection **C**. Rheobase current, the minimum current to elicit an action potential. **D**. Example whole cell current clamp recordings following 200pA current injection. **E**. F-I curve showing the frequency of action potentials elicited by a 50pA stepwise increase in injected current lasting 500msec (mean±se), p=1.26e-10 for Control and FL13 comparison, p=1.94e-13 for Control and FL19 comparison, n=6-14 cells per group, two-way ANOVA with Tukey HSD. **F**. Harmaline injection and onset of tremor induction, student t-test, p-value = 0.04.

### Pcp2CreTetTKO PCs are hyperactive and preferentially sensitive to harmaline

Given the large number of ion channel genes whose expression is altered in the Pcp2CreTetTKO, we were interested in determining whether these changes prevented PCs from attaining their mature electrophysiological properties. To address this issue, patch clamp recordings from PCs in adult, acute slice preparations were performed in both knockout lines (FL13 and FL19) their respective heterozygous knockout (Pcp2CreTetTKO-het) controls, and Pcp2CreTetFlox mice. We found no significant changes in resting membrane potential or input resistance, indicating unaltered intrinsic electrophysiological and morphological properties (Figure 5A-B). While we observed no changes in rheobase current (Figure 5C), a fixed current injection elicited dramatically higher rates of neuronal firing in Pcp2CreTetTKO PCs (Figure 5D). F-I curves demonstrated that homozygous knockouts have significantly higher PC firing rates than Pcp2CreTetTKO-het or Pcp2CreTetFlox across a series of escalating current injections (Figure 5E). Although it is difficult to predict which ion channel alterations might contribute to hyper-excitability as a result of loss of 5hmC, studies of the G protein-activated inwardly rectifying potassium channel Girk1 (Kcnj3) suggest that it may contribute to this phenotype. Girk1 activation decreases PC excitability by inhibiting both sodium and calcium spikes, and it modulates the complex spike response elicited by climbing fiber activation (Lippiello et al, 2020). Thus, the strong decrease in Girk1 expression we have observed in the Pcp2CreTetTKO PC suggests that diminished Girk1 may contribute to enhanced PC excitability evoked by stepwise current injections.

The heightened excitability suggested that the Pcp2CreTetTKO mice might exhibit increased sensitivity to harmaline, a tremorgenic drug whose action elicits abnormal Purkinje cell activity (Brown et al., 2020). To assess this possibility, the sensitivity to harmaline was measured as the time required for tremor onset post injection. As predicted, a shorter onset interval was observed in the Pcp2CreTetTKO animals (Fig. 5F). This demonstrates that the PC hyperexcitability can result in altered function of the cerebellar circuitry and motor behavior, at least in response to induced climbing fiber activation.

## DISCUSSION

Since the discoveries that 5hmC is present at high levels in mammalian neuronal genomes (Kriaucionis and Heintz, 2009) and that it is produced from 5mC by the Tet oxidases (Tahiliani et al., 2009), its possible roles as a stable epigenetic mark and as an intermediate in DNA demethylation have been intensively investigated (Wu and Zhang, 2017). The present study adds to a growing body of work demonstrating that 5hmC plays a critical role in development and function of the nervous system. A central finding of this study is that continued Tet activity and 5hmC accumulation during PC differentiation is necessary for acquisition of their refined functional properties. The consequences of loss of Tet function in PCs include failure to advance transcriptional and epigenetic programs as differentiation continues, and altered electrophysiological and behavioral responses that are typical of mature PCs. It is noteworthy that these phenotypes are evident despite Cre expression loss of Tet function in the Pcp2CreTetTKO late in PC differentiation. It seems likely based on these findings, and the gradual accumulation of 5hmC evident in our data, that there is a continual requirement for 5hmC as a driver for epigenetic remodeling in differentiating, postmitotic neurons. Although studies of neural stem cells (Li et al., 2015; Xu et al., 2012, p. 1), cerebellar granule cells (Zhu et al., 2016) and embryonic brain development have suggested that 5hmC plays a significant role in neural progenitors, a continual requirement for 5hmC may be particularly important for Purkinje cells, pyramidal cells, and other long range projection neurons whose differentiation program includes postmitotic development of sophisticated morphological features and abundant synapses.

A second principle that has emerged from our studies is that Tet mediated active DNA demethylation occurs in postmitotic neurons. Our finding that loss of both 5mC and 5hmC in a specific subset of highly expressed gene bodies and ATAC accessible regulatory sites requires TET activity provides critical in vivo evidence supporting studies in vitro (Weber et al., 2016) and in cultured cells (He et al., 2011) that have defined this pathway. For some genes (Gpr63, Grid2) the progression toward loss of modified C is easily appreciated. In these cases, loss of 5mCG and accumulation of 5hmCG over the gene body is clearly evident between P0 and P7, and continued Tet activity is required to remove both residual 5mCG and 5hmCG to result in DNA demethylation in the adult. Although we have been unable to detect 5fC and 5caC as transient intermediates in this process (Figure S3L), this is not surprising given the very small fraction of the Purkinje cell genome that undergoes active DNA demethylation, and the small amounts of genomic DNA that can be obtained from this very rare cell population. Identification of additional components of the of the DNA demethylation pathway that is occurring in PCs will require further genetic studies disrupting candidate genes in the TDG-BER pathway. Since important non-cell autonomous events are required for PC differentiation (Leto et al., 2016), we believe that additional PC specific knockouts that are restricted to postmitotic cells will be most informative.

An important question arising from these findings is which cells in the nervous system require Tet dependent active DNA demethylation to complete their developmental programs. Although it is probable that other large neurons with complex and prolonged postmitotic differentiation programs share this requirement, we have not yet detected active DNA demethylation in cerebellar granule cells, whose development includes a protracted proliferative phase in which 5hmC can participate in replication dependent DNA demethylation. Furthermore, given reports that remodeling of DNA methylation may occur in response to a wide variety of stimuli in the adult mice (Kaas et al., 2013; Rudenko et al., 2013), additional studies of Pcp2CreTetTKO and other precisely constructed mouse models will help to understand possible functions of 5hmC in neuronal plasticity.

A third advance reported here is the collection of very high-resolution data to understand details of the relationships between DNA methylation and hydroxymethylation, chromatin accessibility and chromatin organization in a single complex neuronal cell type as it develops from a primitive neuronal precursor to a fully articulated adult neuron. These data have highlighted several features of epigenetic development that remain to be explored, including the definition of two distinct relationships between non-CG DNA methylation/hydroxymethylation and transcriptional repression, the mechanism for selection of specific genes as substrates for active DNA demethylation, and the requirement for 5hmC for proper expression of specific genes involved in modulation of PC excitability (e.g. Kcnj3). Although our data reinforce prior studies of the general relationships between DNA methylation, transcription and chromatin structure, we note that there are many exceptions that require further investigation. One intriguing example is the very different relationships between genes and de novo methylation in CG and non-CG contexts (Lister et al., 2013). In Purkinje cells, as in other neurons, substantial enrichment of 5hmCG relative to its general accumulation genome wide occurs in active transcription units, suggesting a mechanism that targets Tet activity to these regions (Figure 2C). While accumulation of 5mCH and 5hmCH can occur within a repressed transcription unit with very nice discrimination between the gene and its surroundings (e.g. Pitpnc1), there are many regions of 5mCH and 5hmCH accumulation that either partially overlap with a gene, or that are present in intergenic regions (e.g. Myo5b/Scarna17/Lipg locus). In all of these regions, however, 5hmC accumulation directly reflects 5mCH levels suggesting 5hmCH localization reflects the pattern of de novo DNA methylation rather than targeted Tet activity. We hope that continued analysis of these datasets will provide a stimulus for additional studies of the mechanisms determining the relative distributions of these important events.

### Concluding Remarks

The data we have reported here advances our understanding of three major functions for 5hmC in the nervous system. Based on extensive studies of the requirements for Tet mediated replication dependent passive DNA demethylation in mESCs, the maturation of the germ line, and developing lymphocyte lineages, it is probable that 5hmC is required to provide accessibility to important regulatory sites as neuronal progenitors exit the cell cycle and begin differentiation. The lack of DNA methylation we have observed at P0 over transcription factor genes (Lhx5, Lhx1, Ldb) that are expressed immediately after PCs exit from the cell cycle suggests that this may result from passive demethylation in dividing progenitors. Further support comes from the observation that DNA demethylation has been observed in comparisons of 5mC and 5hmC levels in the frontal cortex of fetal versus adult mouse and human brains, and the finding that a fraction of these loci retain their methylation status in Tet2^−/−^ mice (Lister et al., 2013). A second function discovered in mature neurons involves stable accumulation of 5hmCG within active genes which helps to reverse the repressive effects of MeCP2 in a process we refer to as functional demethylation (Mellén et al., 2017). The finding that elevated 5hmC remains within genes that are repressed during PC differentiation (Figure 2C) adds to this model the fact that functional demethylation is not sufficient to maintain gene expression. Finally, we report here that Tet mediated active DNA demethylation is required for proper expression of a subset of highly expressed PC specific genes, adding a third important function for 5hmC in the nervous system. These findings place additional emphasis on investigation of ongoing functions of the TDG-BER pathway as elements of normal neuronal development rather that responses to accumulating DNA damage. Given these data, and recent studies linking Tet mutations to neurodegenerative disease (Cochran et al., 2020, p. 2; Marshall et al., 2020, p. 2), it is evident that further exploration of each of these three functions in the context of development, aging and degeneration will continue enhance our understanding of 5hmC and the brain.

## Acknowledgments

We thank all Heintz lab members from the past and present for thoughtful discussions and suggestions. We thank Thomas Carroll, Ji-Dung Luo, Matthew Paul, Doug Barrows and Wei Wang from the Rockefeller University Bioinformatics Resource Core; Svetlana Mazel, Selamawit Tadesse, Stanka Semova, Songyan Han and Samer Shalaby from The Rockefeller University Flow Cytometry Resource Center; Connie Zhao and Christine Lai from The Rockefeller University Genomics Resource Center; Rada Norinsky from the Rockefeller University Transgenic and Reproductive Technology Center.

## Author contributions

E.S. and N.H. designed this study and wrote the manuscript, E.S. and M.R. conducted the experiments. A.R. generated resources and commented on the manuscript.

## Data and materials availability

Pcp2CreTetTKO mice will be made available with a Material Transfer Agreement (MTA). Sequencing data will be available on the NCBI Gene Expression Omnibus upon publication. Code for data analysis will be available on GitHub upon publication.

**Fig. S1.**
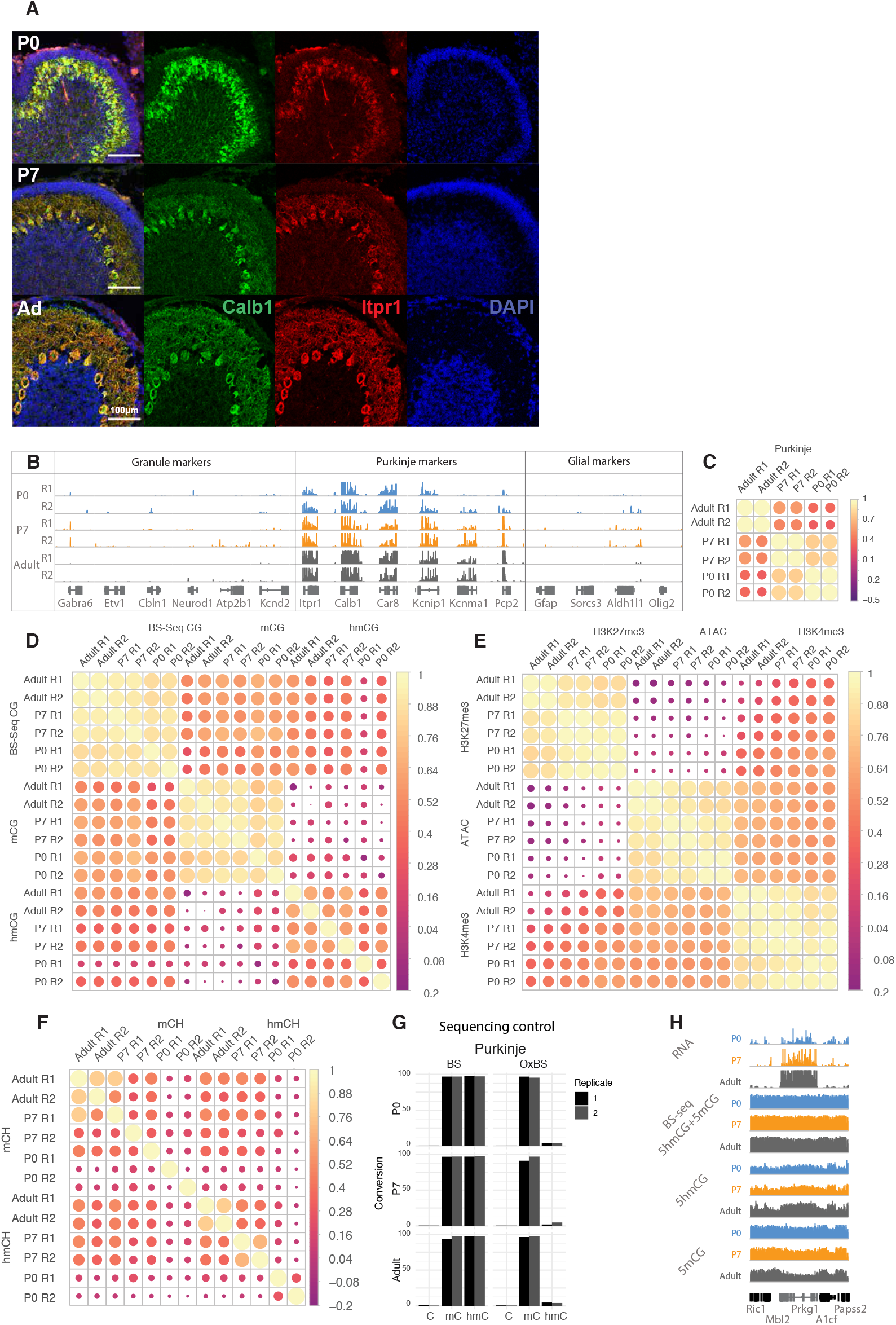
Quality control of sequencing datasets. **A**. Immunofluorescence staining of Purkinje cells at P0, P7 and adult timepoints with Calb1 (green) and Itpr1 (red). **B.** IGV representation of Purkinje specific markers (Itpr1, Calb1, Car8, Kcnip1, Kcnma1, Pcp2) enrichment and depletion of granule (Gabra6, Etv1, Cbln1, Neurod1, Atp2b1, Kcnd2) and glial (Gfap, Sorcs3, Aldh1l1, Olig2) markers. **C**. Pearson correlation of RNA-Seq datasets. **D**. Pearson correlation of gene body accumulation of 5hmCG+5mCG (BSSeq), 5hmCG and 5mCG (derived from maximum likelihood estimator model of BSSeq and OxBSSeq). **E**. Pearson correlation of ATACSeq, H3K4me3 and H3K27me3 ChIPSeq dataset enrichment +/− 5kb around the gene promoters. **F**. Pearson correlation of gene body accumulation of 5hmCH and 5mCH (derived from maximum likelihood estimator model of BSSeq and OxBSeq). **G**. Bisulfite conversion and oxidation efficiency in BSSeq and OxBSSeq datasets. **H**. IGV representation of BS-Seq (showing 5hmCG+5mCG signal together) and OxBS-Seq (showing 5hmCG+5mCG signal separately) at the three developmental timepoints

**Fig. S2.**
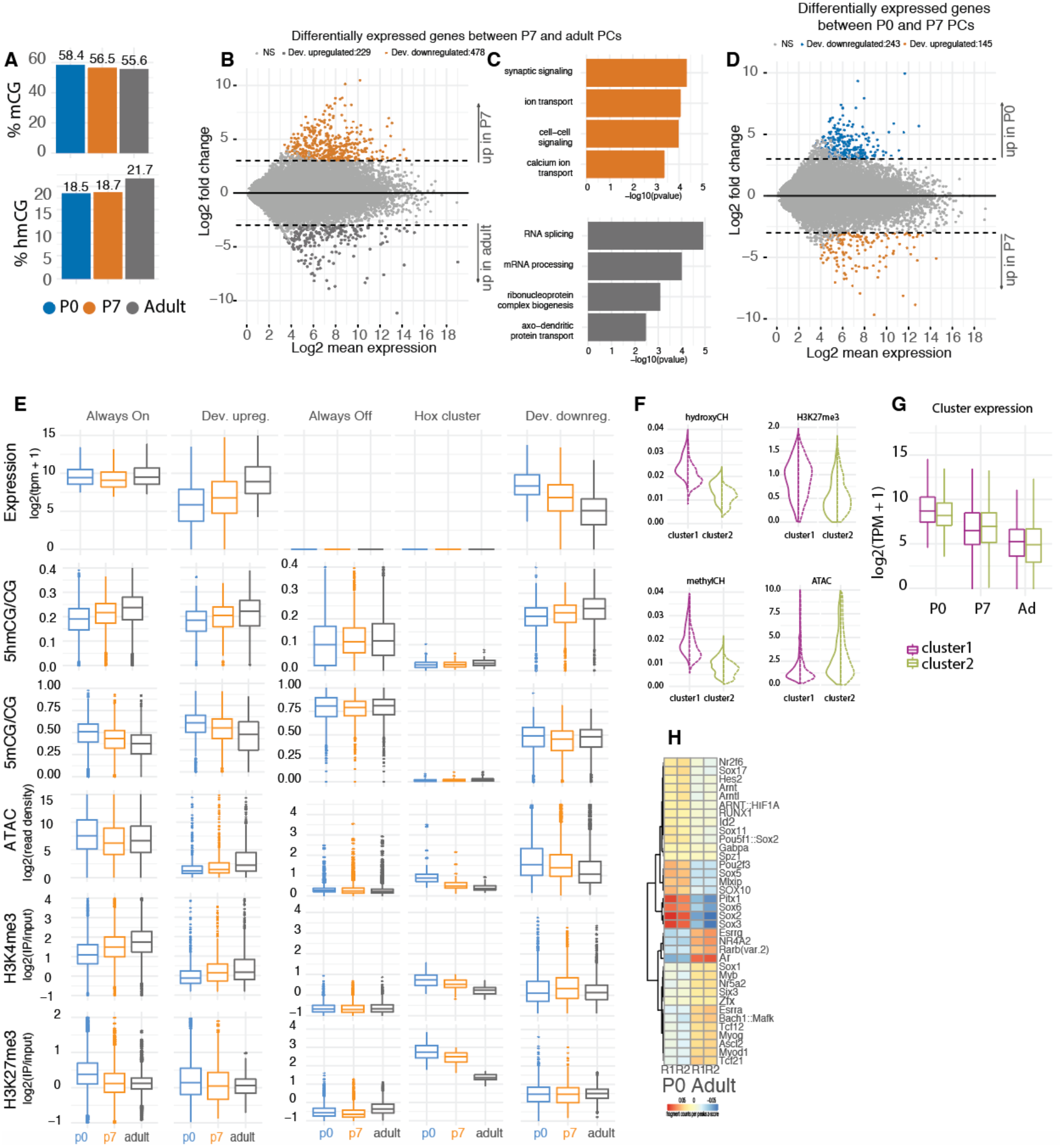
Gene expression and chromatin dynamics during Purkinje cell development. **A**. Genome wide quantification of the levels of 5hmCG and 5mCG at each timepoint. **B**. MA plot representing differential gene expression between P7 and adult. Orange shaded dots represent genes enriched in P7 and gray dots represent genes enriched in adult, with absolute log2 fold change of 2 and p-adj value of 0.05. **C**. GO analysis of differentially expressed genes between P7 and adult. **D**. MA plot representing differential gene expression between P0 and P7 timepoints. Blue shaded dots represent genes enriched in P0 and orange dots represent genes enriched in P7, with absolute log2 fold change of 2 and p-adj value of 0.05. **E**. Quantification of expression, chromatin accessibility, 5mCG, 5hmCG, H3K4me3 and H3K27me3. **F**. Violin plots of the quantifications of 5hmCH, 5mCH, ATAC and H3K27me3 density over the two clusters of repressed genes. **G**. Expression levels log2(TPM+1) of the genes in each cluster. **H**. Differentially abundant motifs of transcription factors in chromatin accessibility data between P0 and Adult PCs.

**Fig. S3.**
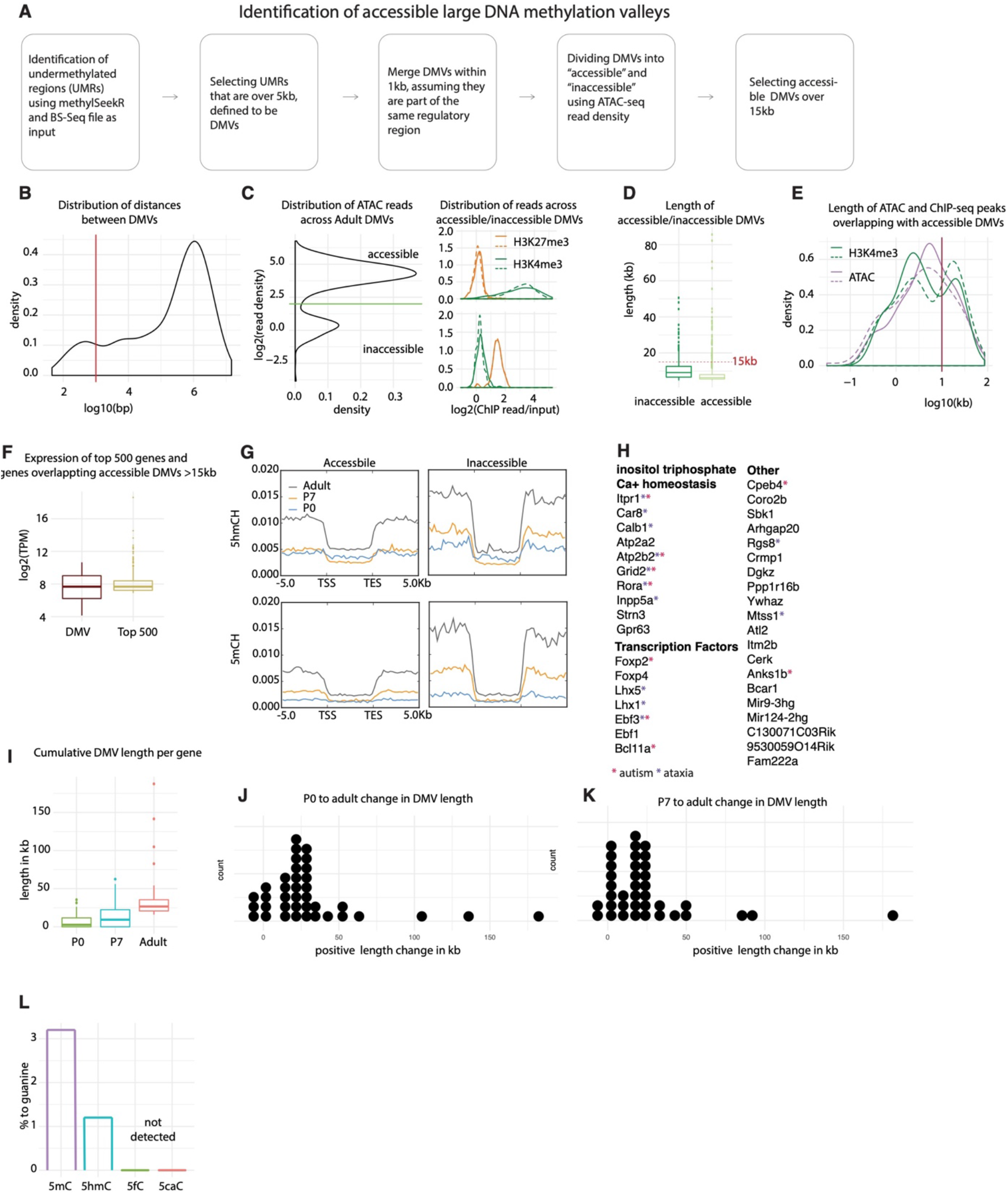
Characterization of Purkinje-specific DNA methylation valleys. **A**. Schematic of DMV identification and filtering. **B**. Distribution of distances between DMVs. DMVs within 1kb of each other are assumed to be part of the same regulatory region and therefore merged. **C**. ATACSeq read density over DMVs shows a bimodal distribution, segregating DMVs into accessible and inaccessible (left panel). Inaccessible DMVs show enriched for H3K27me3 reads, while accessible DMVs show enrichment for H3K4me3 reads (right panel, dotted and straight lines designate replicates). **D**. Distribution of lengths of accessible and inaccessible DMVs. **E**. Length of broad peaks of ATACSeq and H3K4me3 ChIPSeq overlapping large accessible DMVs. **F**. Expression of the top 500 most expressed genes in Purkinje cells and the genes associated with DMVs. **G**. DMVs do not accumulate any modifications in CH context. **H**. Genes associated with DMVs, red star denotes role in autism, gray in ataxia **I**. Cumulative length of DMVs per gene. **J-K**. Change of DMV length between P0 and adult (**J**) and P7 and adult (**K**). **L**. Mass spectrometry analysis of genomic DNA from adult PCs.

**Fig. S4.**
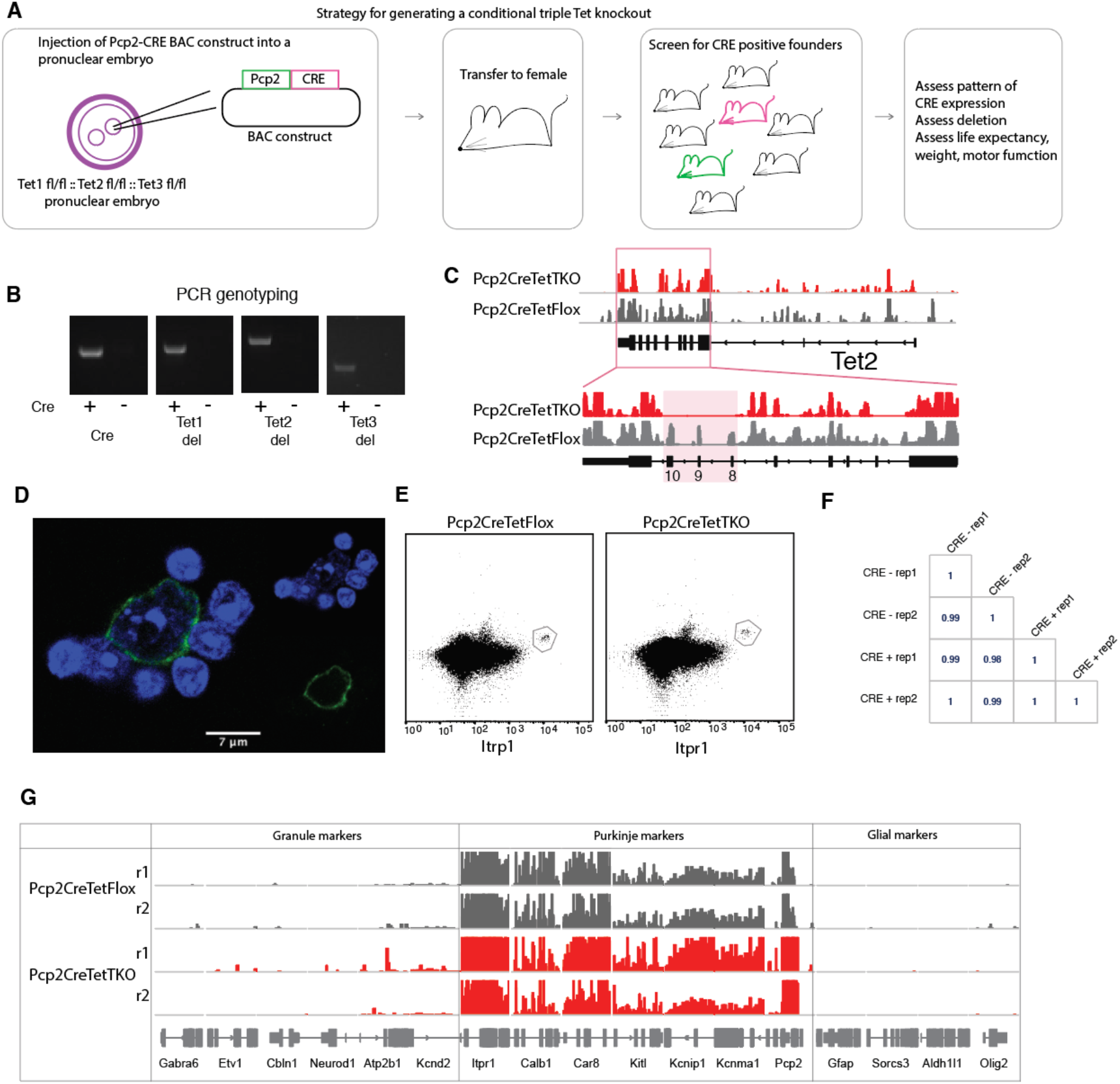
Generation of Pcp2 cre/wt :: Tet1 del/del :: Tet2 del/del :: Tet3 del/del (Pcp2CreTetTKO) mouse founder lines. **A.** Schematic of Pcp2CreTetTKO generation via pronuclear injection. **B**. PCR genotyping for CRE, Tet1 deletion, Tet2 deletion and Tet3 deletion in adult Pcp2CreTetTKO Purkinje genomic DNA. **C**. Deletion of Tet2 exons evident in nuclear RNASeq Pcp2CreTetTKO. **D**. Example of Pcp2CreTetTKO nuclei stained with Itpr1 post dissociation and pre-sorting, counterstained with DAPI, a heterochromatin marker **E**. Example FANS plot of adult Pcp2CreTetTKO nuclei. **F**. Pearson correlation between TKO and control. **G**. IGV representation of Purkinje specific markers (Itpr1, Calb1, Car8, Kitl, Kcnip1, Kcnma1, Pcp2) enrichment and depletion of granule (Gabra6, Etv1, Cbln1, Neurod1, Atp2b1, Kcnd2) and glial (Gfap, Sorcs3, Aldh1l1, Olig2) markers in Pcp2CreTetTKO and Pcp2CreTetFlox.

**Fig. S5.**
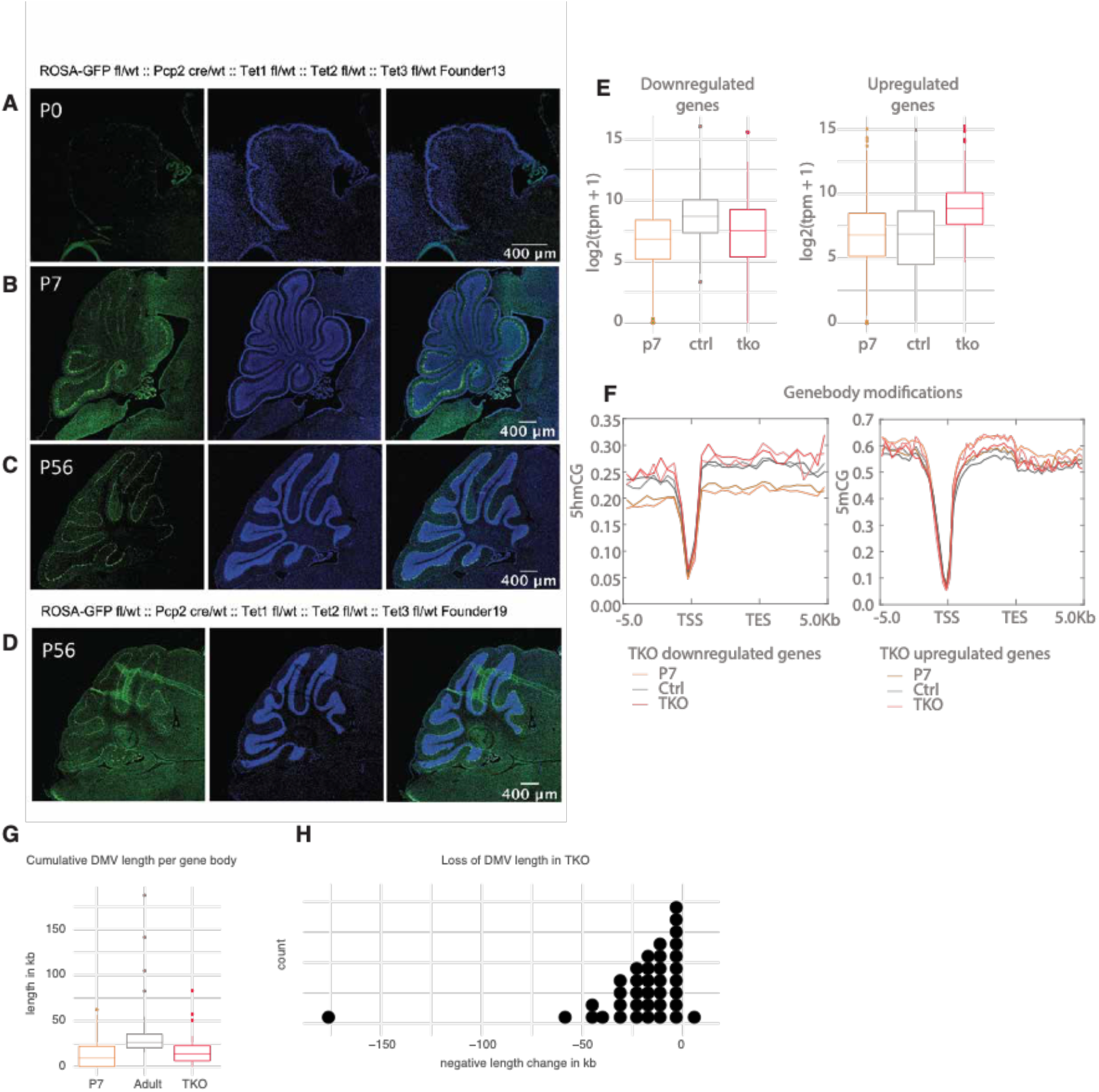
Cytosine dynamics in Pcp2CreTetTKO. **A-C**. Pcp2 cre/wt :: Tet1 del/del :: Tet2 del/del :: Tet3 del/del mice were crossed to ROSA26-eGFP reporter line to evaluate CRE expression during development. Panels represent murine cerebella at P0 (**A**), P7 (**B**) and P56 (**C**) timepoints in FL13, stained with anti-GFP antibody. **D**. anti-GFP staining in P56 FL19. **E**. Expression of genes with increased and decreased expression in Pcp2CreTetTKO (boxplots in left panel, heatmaps in right panel). **F**. Metagene plot of mean mCG and hmCG modifications over promoters and genebodies of genes with increased or decreased expression in Pcp2CreTetTKO. **G**. Cumulative DMV length per gene body between Pcp2CreTetTKO and Pcp2CreTetFlox. **H**. Change in cumulative DMV length between adult Pcp2CreTetTKO and Pcp2CreTetFlox.

**Table S1.**
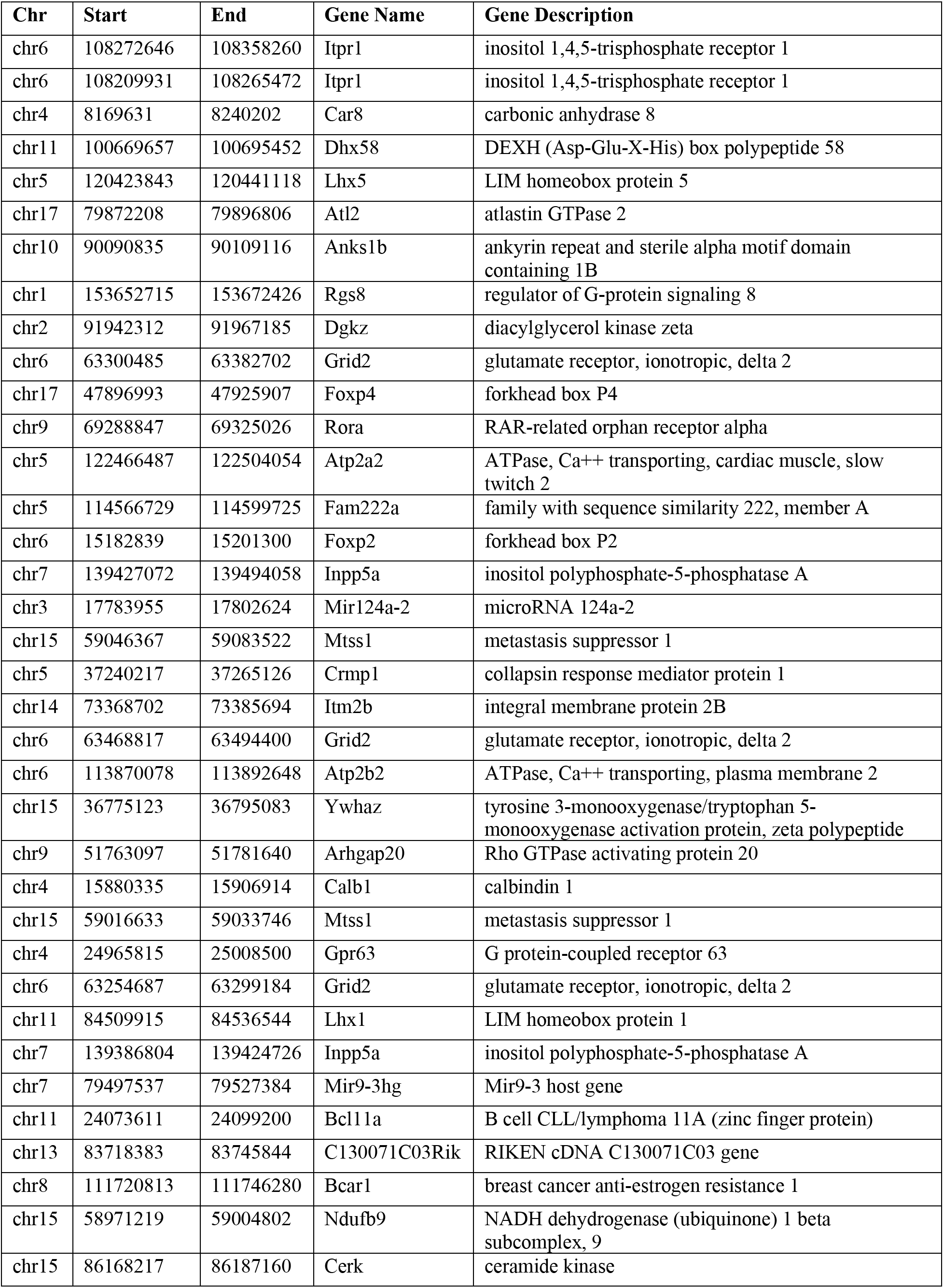
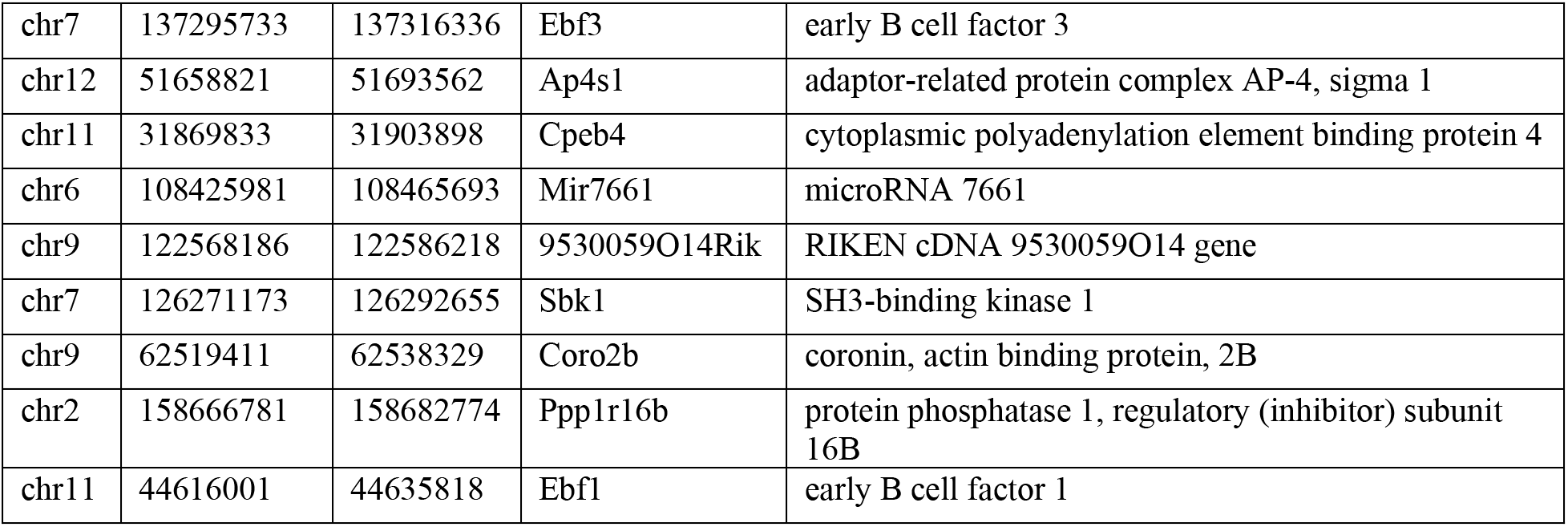
DMVs identified in Adult Purkinje neurons.

## Methods and Materials

### Animals

Wildtype C57BL6/J (RRID:<u>IMSR_JAX:000664</u>) were obtained from Jackson Laboratories. Animals were maintained on a 12hr light/12hr dark cycle with food and water ad libitum. Animal protocols were approved by the Rockefeller University Institutional Animal Care and Use Committee, in accordance with the US National Institutes of Health Guide for the Care and Use of Laboratory Animals.

### Pcp2CreTetTKO murine strain generation

Tet1fl/fl::Tet2fl/fl::Tet3fl/fl animals were a gift from Anjana Rao. We used a previously characterized Pcp2-CRE BAC (RP24-186D18) construct from the GENSAT project (Gong et al., 2007, 2003) to create a conditional TKO in Purkinje cells. Pronuclear injections of the BAC construct into zygotes were performed at the Transgenic and Reproductive Technology Center at the Rockefeller University.

### Cell type specific nuclei isolation

#### Nuclei isolation

Nuclei isolation protocol was followed as described (Xu et al., 2018). Briefly, mouse cerebella were dissected and flash frozen using liquid nitrogen. To isolate nuclei, tissue was thawed on ice for 30 min and then transferred to 5mL of homogenization buffer (0.25 M sucrose, 150 mM KCl, 5 mM MgCl2, 20 mM Tricine pH 7.8, 0.15 mM spermine, 0.5 mM spermidine, EDTA-free protease inhibitor cocktail, 1mM DTT, 20U/mL Superase-In RNase inhibitor, 40U/mL RNasin ribonuclease inhibitor). Tissue were homogenized by 30 strokes of loose (A) followed by 30 strokes of tight (B) glass dounce. Homogenate was supplemented with 5mL of a 50% iodixanol solution (50% Iodixanol/Optiprep, 150 mM KCl, 5 mM MgCl2, 20 mM Tricine pH 7.8, 0.15 mM spermine, 0.5 mM spermidine, EDTA-free protease inhibitor cocktail, 1mM DTT, 20U/mL Superase-In RNase inhibitor, 40U/mL RNasin ribonuclease inhibitor), and laid on a 27% iodixanol cushion. Nuclei were pelleted by centrifugation 30 min, 10,000 rpm, 4°C in swinging bucket rotor (SW41) in a Beckman Coulter XL-70 ultracentrifuge. The nuclear pellet was resuspended in homogenization buffer.

#### Nuclei labeling and sorting

Nuclei were fixed with 1% formaldehyde for 8 minutes at room temperature (RT) with mild agitation. The crosslinking reaction was quenched with 0.125M glycine for 5 minutes at RT. Nuclei were pelleted at 1000g, 4 minutes, 4C, and then washed two times with Wash Buffer (PBS, 0.05% TritonX-100, 50ng/mL BSA, 1mM DTT, 10U/uL Superase-In RNase Inhibitor). Nuclei were blocked with Block Buffer (Wash buffer with an additional 50ng/mL BSA) for 30 minutes at RT, incubated with primary antibody for 1 hour at RT, and then washed three times with Wash Buffer with spins in between washes as described above. Nuclei were then incubated in secondary antibody for 30 minutes at RT and washed three times with Wash Buffer. Primary and secondary antibodies were diluted in Block Buffer. Secondary antibodies were purchased from Life Technologies or Jackson Immunoresearch and were used at 1:500 dilution. Secondary antibodies from goat used: mouse Alexa488, rabbit Alexa594.

#### FACS

Nuclei were stained with DyeCycle Ruby to 20uM final concentration. Nuclei were sorted using a BD FACSAria cell sorter using the 488nm and 561nm lasers. First, samples were gated using DyeCycle Ruby to select only singlets, then appropriate populations were gated based on their separation. Analysis was performed using FlowJo software. Qiagen Buffer PKD was added to the sorted nuclei and the samples were stored at −80C.

### Immunohistochemistry

Mice were decapitated at P0 and P7, and the brains were dissected, and immersion fixed in 4% formaldehyde (w/v) overnight at 4C. Adult mice were deeply anesthetized and then the brains were fixed by transcardiac perfusion with PBS followed by 4% formaldehyde. Brains were further fixed by immersion fixation in 4% formaldehyde overnight at 4C. All brains were cryoprotected in 30% sucrose in PBS, embedded in OCT and cut with a Leica CM3050 S cryostat into 20um sections. The sections were immediately mounted on slides and stored at −20C. Antigen retrieval using sodium citrate buffer (10mM sodium citrate, 0.05% Tween 20, pH 6.0) was performed by heating the slides to 95-100C and then 10-minute incubation in the microwave at the lowest power. The slides were cooled off to room temperature for 1 hour. The slides were washed in PBS and blocked with 3% BSA in PBS with 0.1% TritonX-100 for 30 minutes at room temperature. Primary antibody incubation was performed overnight at room temperature, washed with PBS, incubated with secondary antibody for 1 hour at room temperature, washed with PBS, stained with DAPI (1:10000) for 10 minutes at room temperature and washed three times with PBS. Slides were cover slipped with Prolong Diamond mounting media. Images were acquired using a Zeiss LSM700 confocal microscope using the same acquisition settings for all samples. Further image analysis was done using FIJI.

### Sequencing methods

#### RNA-Seq

RNA and gDNA from fixed nuclei was purified using the Qiagen AllPrep FFPE kit with the following modifications from. After DNA/RNA separation spin, the RNA-containing supernatant was removed and incubated at 65oC for 30 minutes, 70oC for 30 minutes, and 80oC for 15 minutes and then proceeded with the manufacturer’s protocol. gDNA was purified following the rest of the manufacturer’s protocol. RNA quality was determined using Agilent 2100 Bioanalyzer. Purified RNA was converted to cDNA and amplified using the Nugen Ovation RNA-Seq System V2. cDNA was fragmented to an average size of 200bp using a Covaris C2 sonicator (intensity 5, duty cycle 10%, cycles per burst 200, treatment time 120 seconds). Libraries were prepared using the NEBNext Ultra DNA Library Prep Kit for Illumina with NEBNext Multiplex Oligos for Illumina. The quality of the libraries was assessed using the Agilent 2200 TapeStation system with D1000 High Sensitivity ScreenTape. Libraries were sequenced at The Rockefeller University Genomics Resource Center on the Illumina NextSeq 500 to obtain 75bp paired-end reads.

#### ATAC-Seq

ATAC-seq libraries were prepared as described (Buenrostro et al., 2013; Chen et al., 2016) with minor modifications. ~25k fixed nuclei were incubated for 10 minutes in lysis buffer (10 mM Tris pH 7.5, 10 mM NaCl, 3mM MgCl2, 0.1% NP-40). Nuclei were then resuspended in 50 ul 1x TD Buffer containing 2.5 ul Tn5 enzyme from Nextera (Illumina) and incubated for 30 min at 37°C. 200ul of reverse-crosslinking buffer (50 mM Tris–Cl, 1 mM EDTA, 1% SDS, 0.2 M NaCl, 5 ng/ml proteinase K) was added to the samples and they were incubated overnight at 65C with shaking. The samples were then purified using the QiaQuick MinElute columns (Qiagen). Libraries were amplified by PCR using the Q5 High Fidelity Polymerase (NEB) for 12 cycles with barcoded primers. Libraries were size-selected with AMPure XP beads (Beckman Coulter). Libraries were sequenced at The Rockefeller University Genomics Resource Center on Illumina NextSeq 500 to yield 75bp paired-end reads.

#### OxBS-seq

DNA conversion and library preparation were performed using the CEGX TrueMethyl-Seq Whole Genome kit (Cambridge Epigenetix, Cambridge, UK) and after its acquisition from Tecan, the Ultralow Methyl-Seq with TrueMethyl oxBS Module (#0541 and #9513) following the manufacturer’s instructions. Briefly, the DNA was sheared to 800bp using Covaris sonicator and treated with the oxidation agent and bisulfite following the manufacturer’s protocol. Libraries were sequenced on Illumina NextSeq 500 to yield 75 bp paired-end reads. The efficiency of the DNA oxidation and conversion was assessed by interrogating the spike-in Digestion Control and Sequencing Control using the dockerized custom pipeline bsExpress from CEGX (https://bitbucket.org/cegx-bfx/cegx_bsexpress) and fell within the expected ranges (≥90% for hmC conversion in the OxBs reaction, <10% for hmC conversion in the BS reaction (hmC/BS over-conversion error rate), <5% for mC conversion in both the OxBs and BS reactions (over-conversion error rate)).

#### ChIP-seq

Chromatin immunoprecipitation was performed using the Low Cell ChIP-Seq kit (#53048, Active Motif) following the manufacturer’s instructions. Libraries were prepared using the supplied library prep reagents. The samples were sequenced on Illumina NextSeq 500 to yield 75 bp paired-end reads.

### Bioinformatic Data analysis

Most data analysis was done in the R/Bioconductor environment (Huber et al., 2015) in RStudio (https://www.R-project.org/, http://www.rstudio.com/). For general processing, data exploration and visualization we used the tidyverse array of packages, in particular ggplot and dplyr (Wickham et al., 2019).

#### RNA-seq

RNA-seq reads were aligned using STAR (Dobin et al., 2013) and genome assemblies from UCSC. In addition to default STAR parameters, we used the following for paired-end data (-- outFilterMismatchNmax 999 --alignMatesGapMax 1000000 -- outFilterScoreMinOverLread 0 -- outFilterMatchNminOverLread 0 --outFilterMatchNmin 60 -- outFilterMismatchNoverLmax 0.05). Aligned reads were converted to bigwigs for visualization in IGV (Robinson et al., 2011) using deepTools (Ramírez et al., 2016). UCSC gene model annotations for whole genes were downloaded using the UCSC Table Browser tool. Transcript level quantifications was performed using Salmon (v1.1.0) (Patro et al., 2017) and imported for differential expression analysis with tximport (v1.10.1) (Soneson et al., 2015) and DESeq2 (v1.22.2) (Love et al., 2014). For up- and down-regulated genes we selected for a log2 fold change of 2 and p-adjusted value of 0.05. Gene ontology analysis was performed using GOrilla (http://cbl-gorilla.cs.technion.ac.il/) (Eden et al., 2009) or DAVID (https://david.ncifcrf.gov) (Huang et al., 2009).

#### ATAC-seq

Reads were processed with trim_galore (Martin, 2011) with parameters “--stringency 3 -- fastqc --paired”. Trimmed reads were mapped to mm10 using bowtie2 (version 2.1.0) (Langmead and Salzberg, 2012) with parameters “-X 2000 --no-mixed --no-discordant”. Duplicates were removed using samtools (Li et al., 2009). Reads were normalized to RPKM using deepTools bamCompare module, ignoring chrX, chrY, chrM and filtering reads for minimum mapping quality of 30. Metagene and heatmap profiles were generated using deepTools modules computeMatrix, plotProfile and plotHeatmap. Unique fragments under 100 nt were used to call peaks with macs2 (Zhang et al., 2008) with parameters “--nomodel −q 0.01 --call-summits”. Broad peaks were called with macs2 as well with parameters “--nomodel −f BAM --keep-dup all --broad −g mm −B −q 0.01”. The peaks were filtered to remove chrY and chrM peaks and peaks that overlap with the mm10 blacklist from ENCODE (The ENCODE Project Consortium, 2012). DiffBind (v2.10.0) (Ross-Innes et al., 2012) was used to identify differentially accessible chromatin regions using the DESeq2 method. Peaks were selected based on 4-fold difference and q-value of 0.05. Differential motif enrichment was performed using chromVAR (v1.4.1) (Schep et al., 2017).

#### OxBS-seq

Reads were processed with trim_galore with parameters “--stringency 3 --fastqc --paired --clip_R1 5 --clip_R2 10”. Trimmed reads were mapped to mm10 using bismark (Krueger and Andrews, 2011) with parameters “--bowtie2 −p 4 --multicore 4” (v0.20.0). Duplicates were removed using deduplicate_bismark. The following tools from methpipe (v3.4.2) (Song et al., 2013) were used for downstream statistical estimation of the methylation and hydroxymethylation levels. First, bismark aligned reads were converted to the custom .mr format and sorted with default parameters. Methylation calls were extracted with methcounts with default parameters.

Methylation and hydroxymethylation levels were estimated using the mlml tool with default parameters. Genome browser files were generated with bedGraphToBigWig. UMRs and LMRs were identified using MethylSeekR (v1.22.0) (Burger et al., 2013) with m = 0.5 and 5% FDR. DMVs were identified as UMRs ≥5 kb with mean.meth ≤15 and regions within 1kb of each other were merged (bedtools merge −d 1000). mm10 CpG island annotations were downloaded from the UCSC table browser. Unique Large DMVs were selected by excluding regions under 15kb. The regions were annotated using HOMER (Heinz et al., 2010). DMR analysis was performed using methylpy (v1.3.4) (Schultz et al., 2015) with default parameters.

#### ChIP-seq

Reads were processed with trim_galore with parameters “--stringency 3 --fastqc --paired”. Trimmed reads were mapped to mm10 using bowtie2 (v2.1.0) with default parameters. Duplicates were removed using samtools. QC was performed with ChIPQC (Carroll et al., 2014) and that estimated fragment size was used for broad peak calling with macs2 (--nomodel −f BAM --keep-dup all --broad −g mm −B −q 0.01).

### Electrophysiology

At 6-8 weeks of age, mice were deeply anesthetized with ketamine/xylazine (100mg/kg, 10mg/kg b.w.) and perfused with ice cold dissection buffer (2.5 mM KCl, 0.5 mM CaCl_2_·2H_2_0, 7.0 mM MgCl_2_·6H_2_0, 25.0 mM NaHCO3, 1.25 mM NaH_2_PO_4_, 11.6 mM (+)-Sodium L-ascorbate, 3.1 mM sodium pyruvate, 110.0 mM choline chloride, and 25.0 mM glucose. Parasagittal cerebellar sections were collected at a thickness of 300μ on a Microslicer (DTK-1000N) and allowed to recover in artificial cerebrospinal fluid (aCSF) (2.5 mM KCl, 118 mM NaCl, 1.3 mM MgCl_2_2.5 mM CaCl_2,_, 26 mM NaHCO_3_, 1 mM NaH_2_PO_4_, 10 mM glucose) at 32°C for 1 hour. Sections were transferred to room temperature and underwent whole-cell electrophysiological recordings. Dissection buffer and aCSF were kept gassed with carbogen. Slices were transferred to the recording chamber and visualized under an Olympus microscope using a (PCI extended camera). Sections were constantly perfused in aCSF gassed with carbogen, temperature controlled to 32°C (TC-324B). Electrophysiology signals were recorded using a HEKA EPC-10 dual patch clamp amplifier. 4-6MΩ glass capillary pipettes were pulled and filled with internal solution for current clamp (130 K-Gluconate, 5 KCl, 10 HEPES, 2.5 MgCl2. 4 Na2ATP, 0.4 Na3GTP, 10 Na-phosphocretine, 0.6 EGTA) or voltage clamp (115 CsMeSO3, 20 CsCl, 10 HEPES, 2.5 MgCl2, 4 Na2-ATP, 0.4 Na-GTP, 10 Na-phosphocreatine, and 0.6 EGTA). After a GΩ seal was achieved, the membrane was disrupted with short bursts of negative pressure to achieve a whole-cell configuration. Series resistance was measured and ranged between 1 to 20MΩ. In current clamp recordings, a 200ms −50pA hyperpolarizing pulse was used to measure membrane resistance. Rheobase current, the minimal current to elicit an action potential, was determined via stepwise injection of 450pA in 50pA increments. Frequency-current (FI) curves were generated across injected currents and quantified using Axograph (v1.7.6).

### Harmaline injections

8wk old mice were administered 30mg/kg harmaline (Sigma-Aldrich, #H1392) through intraperitoneal injection as described (Brown et al., 2020). Harmaline tremor consistently developed in both WT and TKO mice. Behavior was scored using JWatcher (v1.0).

### Antibody list

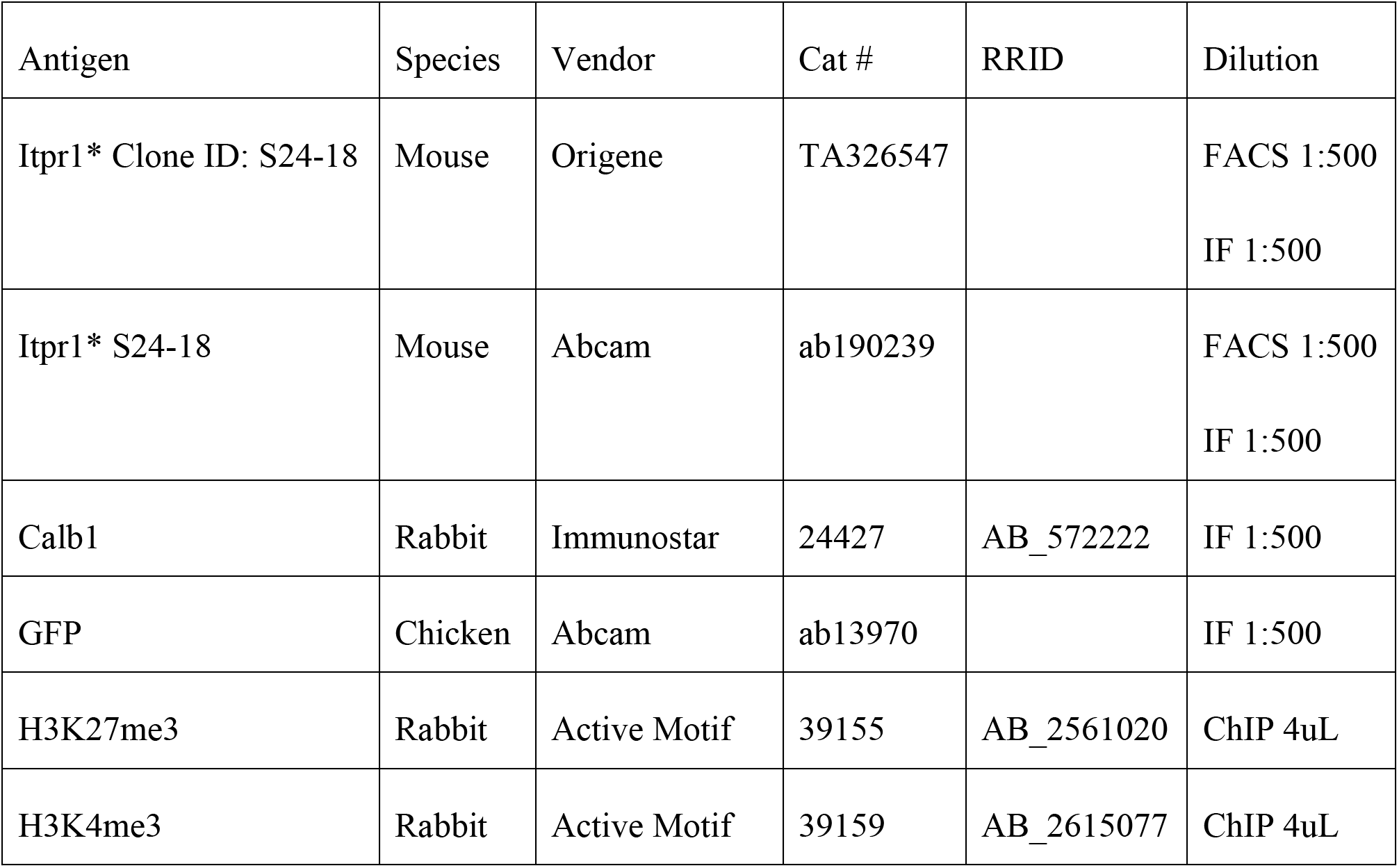
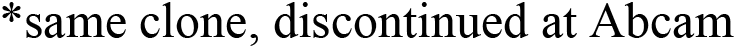

| Antigen | Species | Vendor | Cat # | RRID | Dilution |
| --- | --- | --- | --- | --- | --- |
| Itpr1* Clone ID: S24-18 | Mouse | Origene | TA326547 |  | FACS 1:500<br>IF 1:500 |
| Itpr1* S24-18 | Mouse | Abcam | ab190239 |  | FACS 1:500<br>IF 1:500 |
| Calb1 | Rabbit | Immunostar | 24427 | AB_572222 | IF 1:500 |
| GFP | Chicken | Abcam | ab13970 |  | IF 1:500 |
| H3K27me3 | Rabbit | Active Motif | 39155 | AB_2561020 | ChIP 4uL |
| H3K4me3 | Rabbit | Active Motif | 39159 | AB_2615077 | ChIP 4uL |

\*same clone, discontinued at Abcam

